# Tetraspanner-based nanodomains modulate BAR domain-induced membrane curvature

**DOI:** 10.1101/2022.11.21.517310

**Authors:** Daniel Haase, Christiane Rasch, Ulrike Keller, Annegret Elting, Julia Wittmar, Annette Janning, Martin Kahms, Christian Schuberth, Jürgen Klingauf, Roland Wedlich-Söldner

## Abstract

Topography is a critical feature driving formation and dynamics of protein and lipid domains within biological membranes. The yeast plasma membrane (PM) has provided a powerful model system to study lateral domain formation, including characteristic BAR domain-induced PM furrows. Currently, it is not clear how the components involved in the establishment of these furrows cooperate to precisely regulate local PM topography. Here we report opposing functions for the Sur7 and Nce102 families of tetraspanner proteins in modulating membrane curvature and domain topography. Using STED nanoscopy and freeze-fracture EM we found that Sur7 tetraspanners form multimeric strands at the upper edges of PM furrows, which counteract the forces exerted by BAR domain proteins and prevent membrane tubulation. In contrast, Nce102 tetraspanners are located basal to the Sur7 proteins and promote BAR domain-induced curvature. The segregation of the two tetraspanner-based nanodomains is further supported by differential distribution of ergosterol to the upper edge of furrows and PIP2 lipids at the furrow base. These findings suggest a general role of tetraspanner proteins in sculpting local membrane domains.

## Introduction

As the primary interface between cells and their environment, the plasma membrane (PM) serves a wide range of biological functions including nutrient uptake, metabolic homeostasis, signal transduction and cell-cell communication. In order to perform these tasks, the PM is laterally segregated into a multitude of nanometer or micrometer-sized domains. The formation of these domains has been studied extensively, focusing on either lipid-driven mechanisms such as raft formation (Simons & Ikonen, 1997) and hydrophobic mismatch (Killian, 1998) or protein-based structures such as cortical actin fences (Kusumi *et al*, 2011) and other multimeric networks (Mueller *et al*, 2012). A common theme in this context is the importance of collective and cooperative interactions between large numbers of individual components.

One particularly relevant class of proteins involved in PM organization is made up of the so-called tetraspanners, proteins with four transmembrane domains that often form multimeric complexes. Prominent examples of such assemblies are the tetraspanin webs that organize signaling complexes in immune cells (Boucheix & Rubinstein, 2001; Levy & Shoham, 2005) and the claudin and occludin polymers that form the structural basis for tight junctions in higher eukaryotes (Staehelin, 1974; Lal-Nag & Morin, 2009). Tetraspanner-mediated PM domains are characterized by high spatiotemporal stability, interaction with defined lipids and partner proteins (Levy & Shoham, 2005; Zuidscherwoude *et al*,2015) and by their association with curved membrane regions (Dharan *et al*, 2022; Bari *et al*, 2011).

The budding yeast *Saccharomyces cerevisiae* has been used as a highly informative model for the study of PM domains (Spira *et al*, 2012; Schuberth & Wedlich-Söldner, 2015). Two groups of yeast tetraspanner proteins, the Sur7 (Sur7, Pun1, Fmp45, Ynl194C, Tos7) and Nce102 (Nce102, Fhn1) families, show similarities to mammalian claudins and occludins, respectively. They are prominent components of a yeast PM domain, known as the MCC/eisosome (Malínská *et al*, 2003; Walther *et al*, 2006). MCC/eisosomes are stable furrows, typically 200-400 nm long, 30-50 nm wide and 50-100 nm deep (Strádalová *et al*, 2009; Douglas & Konopka, 2014; Lee *et al*, 2015). The term “MCC” (<u>M</u>embrane <u>C</u>ompartment occupied by <u>C</u>an1) is derived from the association of the arginine permease Can1, whereas “eisosomes” are peripherally associated protein complexes built around two BAR (Bin, Amphiphysin, and Rvs) domain proteins - Pil1 and Lsp1. Together, these proteins assemble into a multimeric coat on the inner surface of the PM in a phosphatidylinositol-4,5-bisphosphate (PIP2) dependent manner (Moreira *et al*, 2009; Karotki *et al*, 2011). This promotes the negative curvature and inward flexure, which gives rise to the characteristic half-pipe shaped furrow that marks this domain as a unique topographic environment within the otherwise flat yeast PM.

MCC/eisosomes and their components have been linked to a variety of biological functions, including protection of nutrient transporters from endocytic internalization (Busto & Wedlich-Söldner, 2019), lipid homeostasis (Young *et al*, 2002; Fröhlich *et al*, 2014) and cell wall synthesis (Alvarez *et al*, 2008; Wang *et al*, 2016; Lanze *et al*, 2020). Multiple links between the local lipid composition and MCC/eisosome function have been proposed. In addition to the role of PIP2 in eisosome assembly (Fröhlich *et al*, 2014), ergosterol has been shown to be enriched within MCC/eisosomes and several eisosomal components such as Nce102, Pkh1/2 and Slm1/2 have been suggested to regulate sphingolipid homeostasis (Walther *et al*, 2006; Grossmann *et al*, 2007; Luo *et al*, 2008; Fröhlich *et al*,2009, 2014; Aguilar *et al*, 2010).

Like many other BAR domain proteins (Frost *et al*, 2009), purified Pil1 and Lsp1 are known to self-assemble into helical structures that induce formation of tubes with diameters of 30 - 40 nm from artificial membranes (Karotki *et al*, 2011). How cells modulate MCC/eisosomal activity to generate stable PM furrows rather than tubes remains unclear (Lanze *et al*, 2020).

In this study we demonstrate that the Sur7 and Nce102 tetraspanner proteins perform distinct and partially opposing functions in modulating the topography of the yeast PM. Using combined stimulated emission depletion (STED) microscopy (Hell & Wichmann, 1994) and freeze-fracture electron microscopy (EM), we show that Sur7 family proteins form multimeric strands on the upper edges of MCC/eisosome furrows, which counteract the forces exerted by BAR domain proteins at the furrow base, and thus prevent membrane tubulation. In contrast, Nce102 tetraspanners are located at the bottom of furrows and promote BAR domain-induced curvature. In addition, the spatial separation of nanometer-scale tetraspanner domains within PM furrows is associated with distinctive lipid environments. While ergosterol is enriched at their Sur7-defined upper edges, our work together with previous reports indicates that PIP2 is concentrated by the BAR domains at the base of furrows. These findings suggest that the local topography of the yeast PM results from an intricate interplay between BAR domain-containing proteins and specific tetraspanner families.

## Results

### Spatial organization of tetraspanner-rich domains in the yeast PM

To characterize the detailed organization of the MCC/eisosome domains we initially made use of two-color total internal reflection fluorescence microscopy (TIRFM) to visualize all domain-resident tetraspanners, and compared their distributions to that of Pil1. The Nce102 paralog Fhn1 was not expressed under our growth conditions and was therefore not included. As additional control we also determined localization for the Sur7 orthologue A08184g from *Kluyveromyces lactis*. All tetraspanners co-localized with Pil1, either when expressed at endogenous levels (Fig EV1A, B) or when overexpressed from the Pma1 promoter (Fig 1A, B). While most Sur7 tetraspanners were strongly concentrated in MCC/eisosomes, Pun1 and Tos7 were also detected throughout the PM irrespective of their expression level, as reflected by the larger network factors (Fig 1C, EV1C). This parameter describes the density of signal distribution for PM proteins (Spira *et al*, 2012). In contrast, Nce102 was exclusively concentrated in MCC/eisosomes under normal conditions (Fig EV1C) but its distribution became more dispersed when overexpressed (Fig 1C). Therefore, labeling of Nce102 was performed exclusively on the endogenous level in all further experiments, whereas expression of Sur7 tetraspanners could be modulated according to the needs to ensure robust comparability of our analysis.

**Figure 1.**
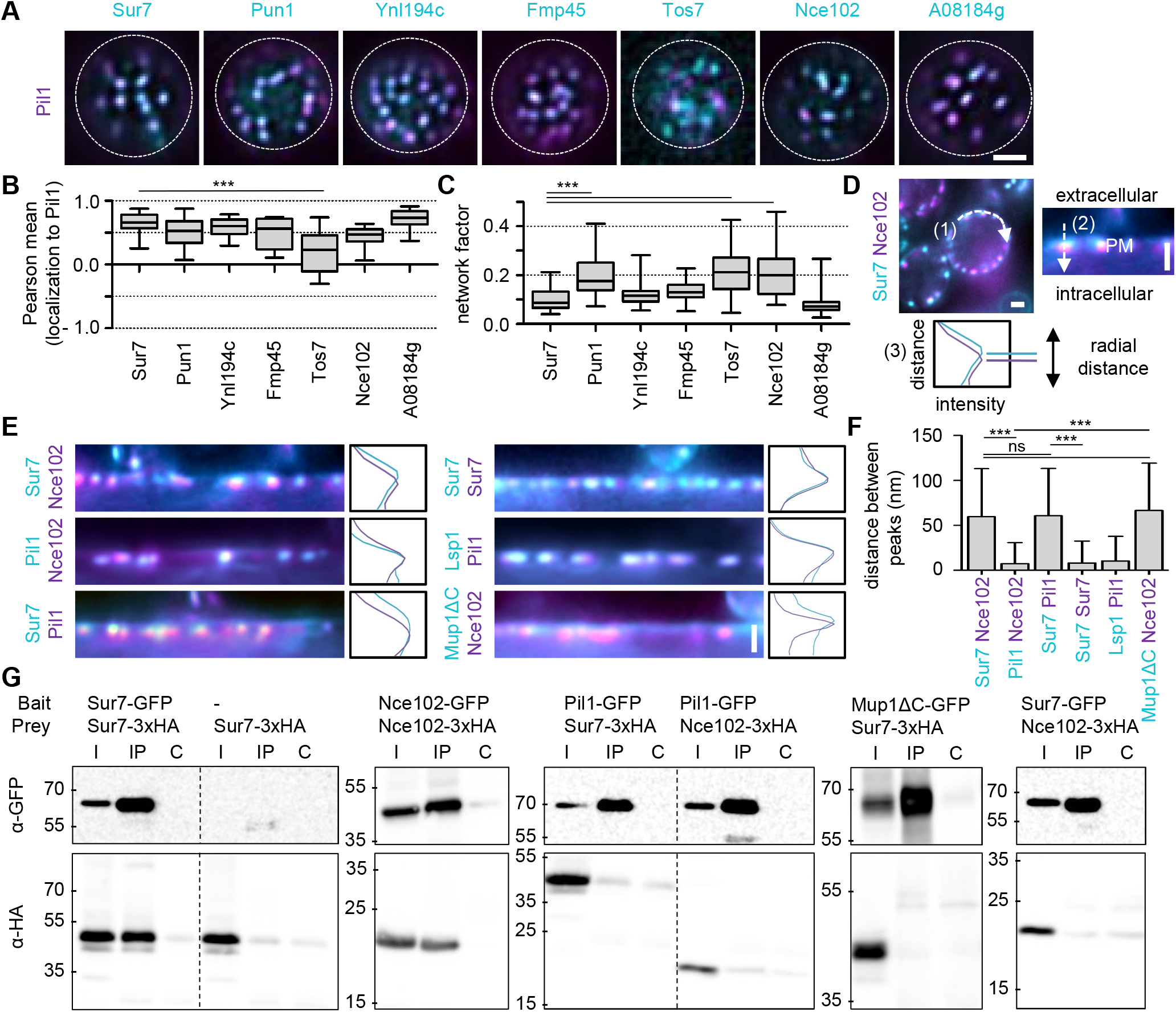
Yeast tetraspanners define subdomains in MCC/eisosomes. A TIRFM images showing colocalization of the indicated tetraspanners (C-terminally fused to GFP (cyan) and expressed from the Pma1 promoter) together with the MCC/eisosomes marker Pil1-RFP (magenta). B Pearson correlation coefficients for colocalization shown in (A). C Lateral distribution of tetraspanners from (A) quantified using the network factor. D Representative linearized profiles were generated along the cell periphery (1) for presentation purposes only. An exemplary two-color intensity profile (3) was taken perpendicularly through a single MCC/eisosome, indicated by a dashed arrow (2). Distances between intensity peaks correspond to radial distances between indicated proteins. E Radial fluorescence intensity distributions of MCC/eisosomal protein pairs and representative intensity profiles perpendicular to MCC/eisosomes. Proteins were endogenously fused at their C-termini to mNeGr (cyan) or mRFPruby (magenta). Note that the linearized profile for Sur7-mNeGr / Nce102-RFP is taken from the same cell as in (D). F Distances of peaks in radial distributions between indicated proteins shown in (E). G Co-Immunoprecipitation of tetraspanners with various targets. Indicated GFP-tagged proteins (bait) were pulled down with Anti-GFP. GFP-tagged bait proteins and HA-tagged prey proteins were detected by Western blot. I: Input, IP: Co-IP with anti-GFP, C: IP with unspecific IgG. Data information: (B, C) Boxplots, ANOVA with Dunnett’s multiple comparison test, n = 15–32 cells (B), n = 41-88 cells (C). (E) Bar graph, ANOVA with Tukey’s multiple comparison test, n > 100 MCC/eisosomes. Scale bars: 1 μm.

To determine the positions of tetraspanners within the MCC/eisosome furrow more precisely, we performed high-resolution radial distance measurements of medial cell sections derived from conventional epifluorescence data (Fig 1D). Using this method, we found that members of the Sur7 family were located approximately equidistant (about 50 nm away) from Nce102, Pil1 and Lsp1 (Fig 1E, F, EV1D). This likely reflected localization at different furrow depths, with Sur7 proteins being closest to the surface. As a representative member of the MCC/eisosome associated symporters, the methionine permease Mup1 exhibited a similar radial distribution to that of Sur7 proteins (Fig 1E, F, depicted is the distribution of a C-terminal deletion variant of Mup1, Mup1ΔC, which exhibits improved localization within MCC/eisosomes, (Busto *et al*, 2018)).

To test whether the apparent colocalization of individual markers might reflect physical interaction between them, we performed co-immunoprecipitation experiments. We found strong interactions among the Sur7 family members (Fig EV1F) and of Nce102 with itself (Fig 1G). Notably, despite their close associations, we found no evidence for physical interactions between Nce102 and Pil1 or Mup1 and Sur7 (Fig 1G).

### Tetraspanners and BAR domain proteins localize to distinct MCC/eisosome subdomains

While we could observe significant and robust separation of two distinct subdomains in radial representations of MCC/eisosomes, the precise relationship between these domains was obscured by the limited resolution of conventional fluorescence microscopes. To improve visualization of subdomains we therefore turned to super-resolution STED microscopy and freeze-fracture EM.

We initially inspected surface sections of cells to determine the lateral localization profiles of key marker proteins for MCC/eisosomes expressed at endogenous levels. The two markers that localized in a more central radial position, Pil1 and Nce102, were found to form single linear strands of around 300 nm in length (Fig 2A). In contrast, the peripherally located Sur7 and Mup1 exhibited a characteristic double-strand appearance (Fig 2A) with 300 nm length and intra-strand gaps of more than 60 nm (Fig 2B). These values were strikingly similar to width and length of MCC/eisosomes observed in EM of freeze-fracture replicas (Fig 2B). Overexpressed Sur7 family members exhibited identical double-strand patterns (Fig EV2A). By combining STED microscopy images of Sur7-HaloTag (Halo) with confocal images of Pil1-mNeonGreen (mNeGr) we confirmed that Sur7 strands laterally overlapped with Pil1-labeled structures and thus indeed represented the Sur7 distribution in MCC/eisosomes (Fig EV2B).

**Figure 2.**
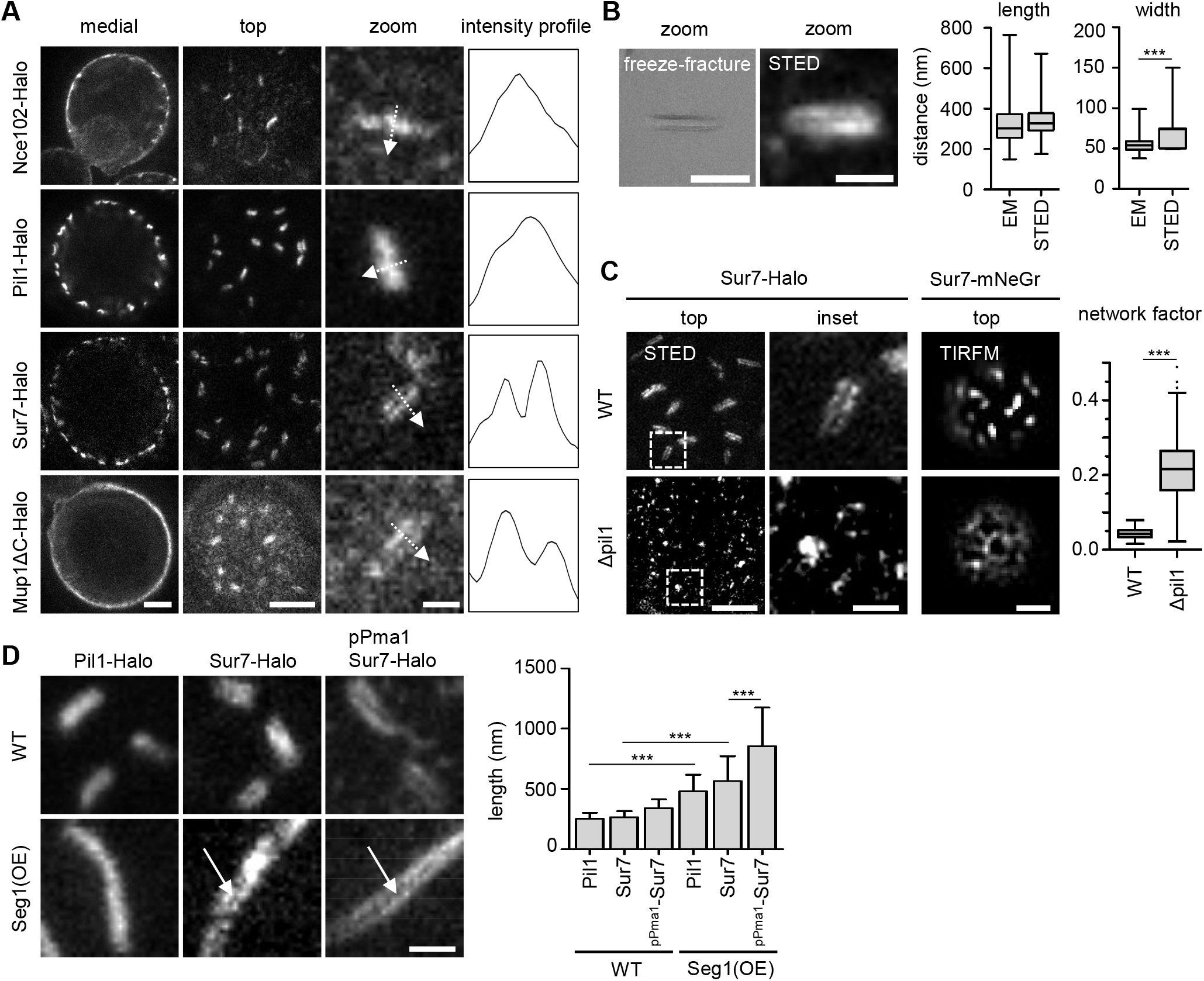
Super-resolution imaging of MCC/eisosome subdomains. A STED microscopy of indicated proteins fused to Halo. Intensity profiles correspond to the dotted lines in the zoomed images. B Comparison of MCC/eisosome furrow dimensions obtained from freeze-fracture (EM) replicas with STED images of a yeast strain expressing Sur7-Halo under the Pma1 promoter. C Representative STED- and TIRFM-images of yeast WT and Δ*pil1* cells expressing Sur7-Halo or Sur7-mNeGr; the latter was quantified in terms of the network factor. D Representative STED images of WT and Seg1 overexpressing (OE) yeast cells expressing either endogenous Pil1-Halo or Sur7-Halo or overexpressing Sur7-Halo from the Pma1 promoter. Arrows highlight double-stand appearance of Sur7-tetraspanners. Quantification shows MCC/eisosomal length upon Seg1 overexpression. Data information: (B, C) Box plots, unpaired t-tests, n > 150 MCC/eisosomes. (D): Bar graph, ANOVA with Tukey’s multiple comparison test, n > 90 MCC/eisosomes. Scale bars: 1 μm, 200 nm (zoom).

As shown previously (Walther *et al*, 2006), deletion of Pil1 led to a complete loss of MCC/eisosome-related structures. In addition to the previously reported eisosomal remnants (Walther *et al*, 2006), Sur7 was distributed over the whole PM and formed small clusters of less than 100 nm diameter (Fig 2C). This was also apparent from an increased network factor in TIRFM images (Fig 2C). On the other hand, overexpression of the eisosomal regulator Seg1, which is involved in the initiation of eisosome formation (Moreira *et al*, 2012), led to the formation of much longer structures that retained the characteristic appearance of the respective markers (Fig 2D). Elongated Sur7 strands in Seg1 overexpressing cells (Seg1OE) remained continuous along both lateral surfaces (Fig 2D), similar to the appearance in WT MCC/eisosomes (Fig 2A, EV2A). Strand separation was easier to distinguish upon Sur7 overexpression, but also led to an increase in Sur7 strand length (Fig 2D).

Taken together, our analysis combining regular epifluorescence, TIRF, STED and freeze-fracture electron microscopy demonstrate that MCC/eisosomes exhibit lateral and radial separation into nanometer scaled domains that are defined by distinct tetraspanner families and BAR domain proteins.

### Sur7 tetraspanners modulate BAR domain-induced PM topography in MCC/eisosomes

Having established the subdomain organization of MCC/eisosomes, we turned to the role of Sur7 tetraspanners and of the parallel Sur7 strands in shaping the local PM topography. We generated a yeast strain lacking all five of the Sur7 family members previously identified in MCC/eisosomes (Fig 1). Strikingly, in this 5xΔ mutant (Δ*sur7*, Δ*pun1*, Δ*fmp45*, Δ*ynl194c*, Δ*tos7*), Pil1 localized to distinctive tube-like structures that remained connected to the PM at their ends (Fig 3A). Importantly, while individual curved structures had track lengths of around 600 nm, their end-end distance of around 250 nm was comparable to that of normal MCC/eisosome furrows (Fig 3B). The tube-like structures correlated with higher Pil1 fluorescence intensities accompanied by a reduction in the number of structures (Fig 3C), indicating that Pil1 is redistributed into fewer but longer structures. An overall conservation of Pil1, Lsp1 and Nce102 levels was confirmed by comparison of protein expression levels (Fig EV3A). To clarify the nature of Pil1-labeled tube-like structures we overexpressed Seg1 (Fig 2D) in the 5xΔ background. We observed pronounced tube-like structures that remained connected to the PM at their ends (Fig EV3B). Importantly, the end-end distance of these structures was comparable to the increased furrow-length of MCC/eisosomes in WT cells overexpressing Seg1 (Fig EV3C), indicating that tube-like structures originate from regular MCC/eisosomes. We then looked at the localization of other MCC/eisosomal makers. Both, Lsp1 and Nce102 were clearly detected within the tube-like structures (Fig 3D), while Mup1 was largely excluded (Fig 3D). We confirmed co-localization of Nce102 and Pil1 within tube-like structures by epifluorescence imaging (Fig 3E) and showed that tube formation depended on the presence of either component (Fig 3F).

**Figure 3.**
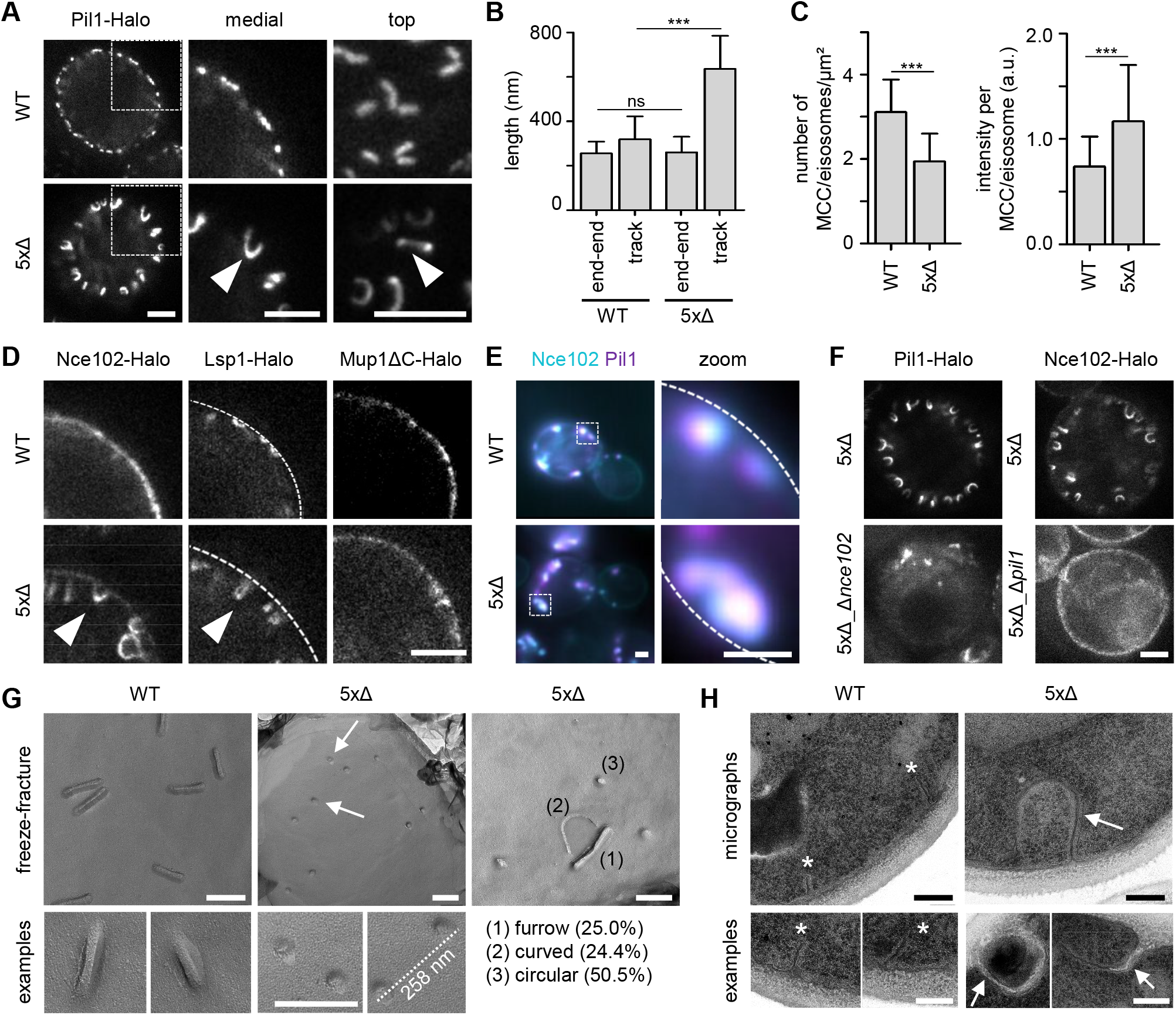
Sur7-tetraspanners prevent closure of MCC/eisosome furrows. A STED images of yeast cells expressing Pil1-Halo in control (WT) and in mutant cells depleted of all five Sur7 tetraspanners (5xΔ). Arrow heads indicate tubular invagination. B Quantification of end-end distance (from top views) or track-length (from medial views) of Pil1-Halo marked structures in yeast WT or 5xΔ cells. C Density (number per area) and intensity of MCC/eisosomes marked by Pil1-Halo. D STED images of yeast WT and 5xΔ cells expressing indicated Halo fusions. Arrow heads highlight tubular structures in 5xΔ cells. Dashed line indicates location of PM. E Colocalization of Pil1-mRFPruby and Nce102-mNeGr in tubular structures of 5xΔ cells. F STED images of 5xΔ and 6xΔ (5xΔ + Δ*pil1* or Δ*nce102*) cells expressing indicated Halo fusions. G Freeze-fracture images of the PM in WT and 5xΔ cells. Arrow indicates likely end of a tubular invagination. The distance between two exemplary circular structures and percentage of identified structural categories (1-3) in 5xΔ cells is shown. H Transmission EM micrographs of ultrathin sections of WT and 5xΔ cells. Asterisks indicate PM furrows and arrows indicate tubular invaginations. Data information: (B, C) Bar graphs, unpaired t-test, n > 44 cells. Scale bars: 1 μm and 200 nm (G, H).

To verify whether the observed structures indeed corresponded to closed membrane tubes, we characterized the ultrastructure of the yeast PM. Freeze-fracture replicas showed the typical 300 nm long furrows of MCC/eisosomes in the WT PM (Fig 3G). We consistently observed different degrees of curvature at the lateral edges vs. furrow tips, as indicated by different levels of contrast (Fig 3G, examples). In the PM of 5xΔ cells we observed a smaller amount of straight furrows (~25%), an equal number of curved invaginations (~25%) and a majority of smaller circular structures (~50%) that fit the expected diameter range seen for normal MCC/eisosomes (Stradalova *et al*, 2009) and Pil1-derived membrane tubes in vitro (Karotki *et al*, 2011). Importantly the distance between neighboring circular structures was comparable to the end-end distance for tubes observed via STED microscopy (Fig 3G, arrows, Fig 3B). Finally, we confirmed the formation of membrane tubes in 5xΔ cells in EM micrographs of 60 nm ultrathin sections. We could identify very distinctive membrane tubes that typically extended over 300 nm into the cell (Fig 3H). Some of these tubes were filled with electron translucent material that was reminiscent in texture to cell wall material (Fig 3H, arrow) and was also seen in Δ*sur7* cells of *C. albicans* (Alvarez *et al*, 2008). This was especially apparent in Seg1 overexpression conditions where tubes were much longer and often twisted (Fig EV3D). These results indicate that Sur7 tetraspanners play a major role in modulating PM topography by preventing the fusion of the upper edges of MCC/eisosome furrows to form closed membrane tubes.

### The function of Sur7 in shaping MCC/eisosomes requires its C-terminal region

To identify specific structural features in Sur7 tetraspanners that mediate their function in PM domain formation, we attempted to rescue the tubulation phenotype by overexpressing individual Sur7 family members in the 5xΔ strain (Fig 4A). We found that three of these - Sur7, Fmp45 and Ynl194c - fully rescued the radial separation between Nce102 and the respective Sur7 tetraspanner (Fig 4A). The single *K. lactis* Sur7 homologue A08184g was also able to fully rescue MCC/eisosome tubulation (Fig 4A). In contrast, the remaining members, Pun1 and Tos7 only led to a partial rescue. (Fig 4A). Notably, these two proteins were the only two Sur7 variants that also localized to PM regions outside of MCC/eisosomes (Fig 1A, C; EV1A, C). The largest differences between sequences of rescuing and non-rescuing Sur7 family members are found in their C-terminal regions (Fig 4B). We therefore repeated our rescue experiments with different Sur7 variants lacking N- and/or C-terminal regions. While deletion of the N-terminus had no obvious effect on the rescue of tubulation, removal of the C-terminus (or removal of both termini) led to a complete loss of rescue activity (Fig 4C). We further confirmed these results by STED microscopy where expression of Sur7 lacking the C-terminus could not prevent formation of tubes in the 5xΔ mutant (Fig 4D). These results are consistent with recent findings in *C. albicans*, which assigned a key function in regulating morphogenesis and stress responses to the Sur7 C-terminus (Lanze *et al*, 2021). Importantly, despite its role in sculpting membranes, the Sur7 C-terminus was not required for recruitment to MCC/eisosomes or for strand formation per se (STED, Fig 4E). Finally, we also confirmed by Co-IP analysis that (like Sur7) Sur7ΔC was able to interact with itself (Fig 4F).

**Figure 4.**
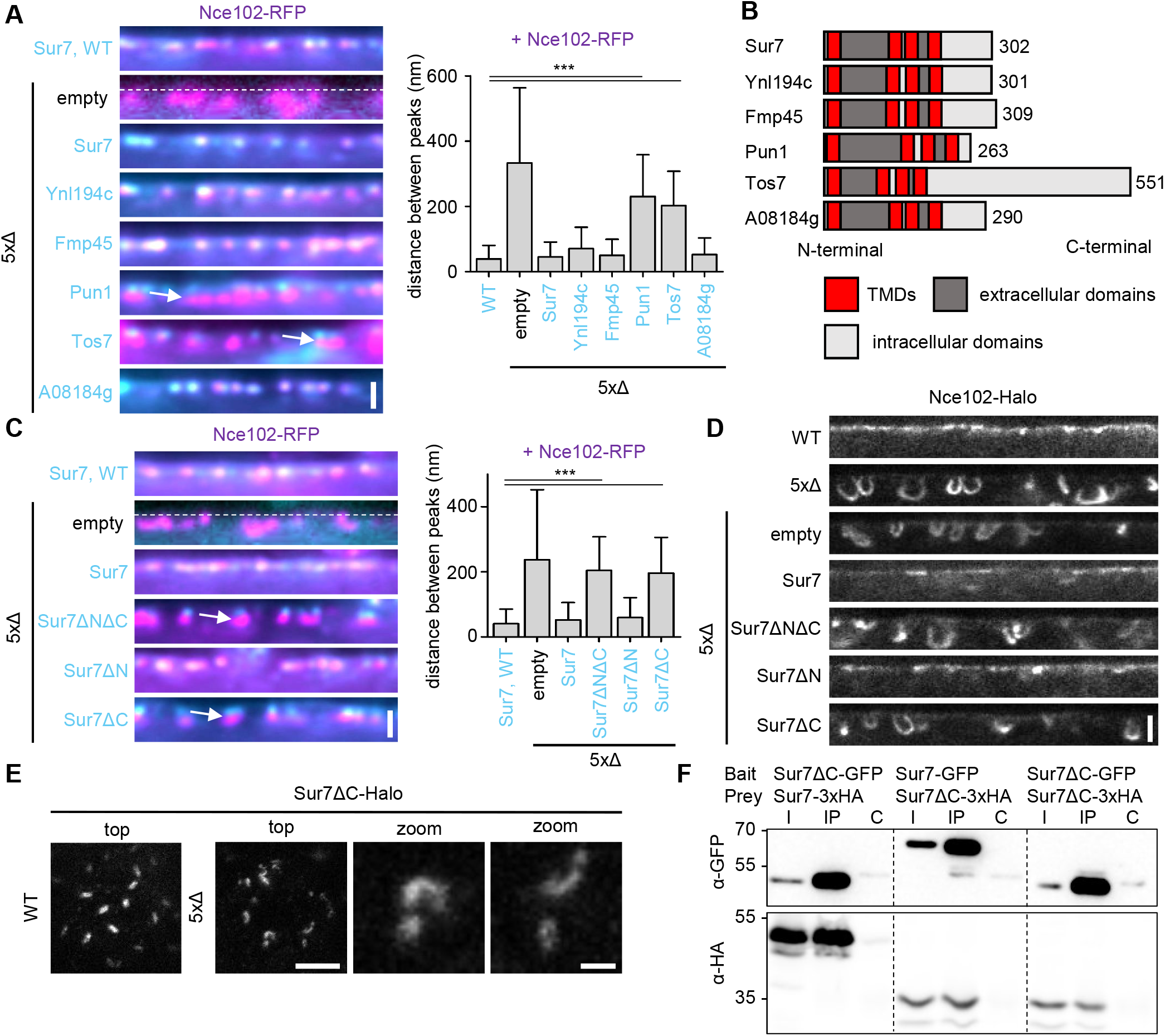
The C-terminus of Sur7 is required for regulating PM domain topography. A Radial fluorescence intensity distributions of indicated protein pairs in linearized profiles of WT and 5xΔ cells expressing Nce102-RFP (magenta) and GFP-fused Sur7-tetraspanners (cyan) from the Pma1 promoter. Bar graph indicates distances between radial profile peaks Arrows indicate separation of tetraspanners. Dashed line indicates location of PM in control. B Topology of Sur7-tetraspanner family members with transmembrane domains (TMDs, red), extracellular (dark grey) and intracellular (light grey) segments indicated. C Radial fluorescence intensity distributions of protein pairs in WT and 5xΔ cells expressing Nce102-RFP (magenta) and GFP-fused Sur7-truncations (cyan) from the Pma1 promoter. Bar graph indicates distances between radial profile peaks. Sur7 = 1-302aa; Sur7ΔC = 1-210aa; Sur7ΔN = 7-302aa; Sur7ΔNΔC = 7-210aa. Arrows indicate separation of tetraspanners. D Linearized STED profiles from 5xΔ cells expressing Nce102-Halo and Sur7 truncations. E STED images of WT and 5xΔ cells expressing Sur7ΔC-Halo from the Pma1 promoter. F Co-Immunoprecipitation of Sur7 truncations. Indicated GFP-tagged proteins (bait) were pulled down with Anti-GFP. GFP-tagged prey proteins and HA-tagged prey proteins were detected by Western blot. I: Input, IP: Co-IP with anti-GFP, C: IP with unspecific IgG. Data information: (A, C) Bar graphs, ANOVA with Dunnett’s multiple comparison test, n > 57 MCC/eisosomes. Scale bars: 1 μm, 500 nm (D), 200 nm (E, zoom).

In summary, the C-terminal cytosolic segment of Sur7 plays a key part in the function of this tetraspanner in counteracting membrane tubulation via the BAR domains of Pil1 and Lsp1.

### Nce102 plays a structural role for MCC/eisosome topography

Considering the critical function of Sur7 tetraspanners in MCC/eisosome morphogenesis, we asked whether the Nce102 tetraspanners have an equally relevant function in PM domain formation. We found that deletion of Nce102 led to the previously reported drastic reduction of MCC/eisosome number, which could be rescued by expression of the non-phosphorylatable Pil1 variant Pil1-4A (Fig 5A), (Walther *et al*, 2007). The radial position of Pil1 was not significantly altered upon deletion of Nce102 or expression of the Pil1-4A variant alone (Fig 5B). However, we found a slightly reduced distance between Pil1-4A and Sur7 in the absence of Nce102 (Fig 5B). Strikingly, in the remaining MCC/eisosomes formed in Nce102-depleted cells, the distance between the two Sur7 strands was markedly increased (Fig 5C). By averaging the Sur7 signal from multiple aligned structures we were able to highlight their parallel stranded appearance (Fig 5D). We next combined the 5xΔ strain with the deletion of Nce102. The resulting cells exhibited only very small number of Pil1-positive structures and these were then often clustered (Fig 5E), which precluded robust quantifications (Fig 5F). However, we could consistently observe Pil1-positive tubes in the 6xΔ deletion strain, which was rescued by Pil1-4A expression (Fig 5E). However, the tubes in 6xΔ cells expressing Pil1-4A were significantly shorter than those found in the 5xΔ control cells (Fig 5F). These results indicate that Nce102 plays a role in MCC/eisosome topography that extends beyond its regulatory effect on Pil1 phosphorylation.

**Figure 5.**
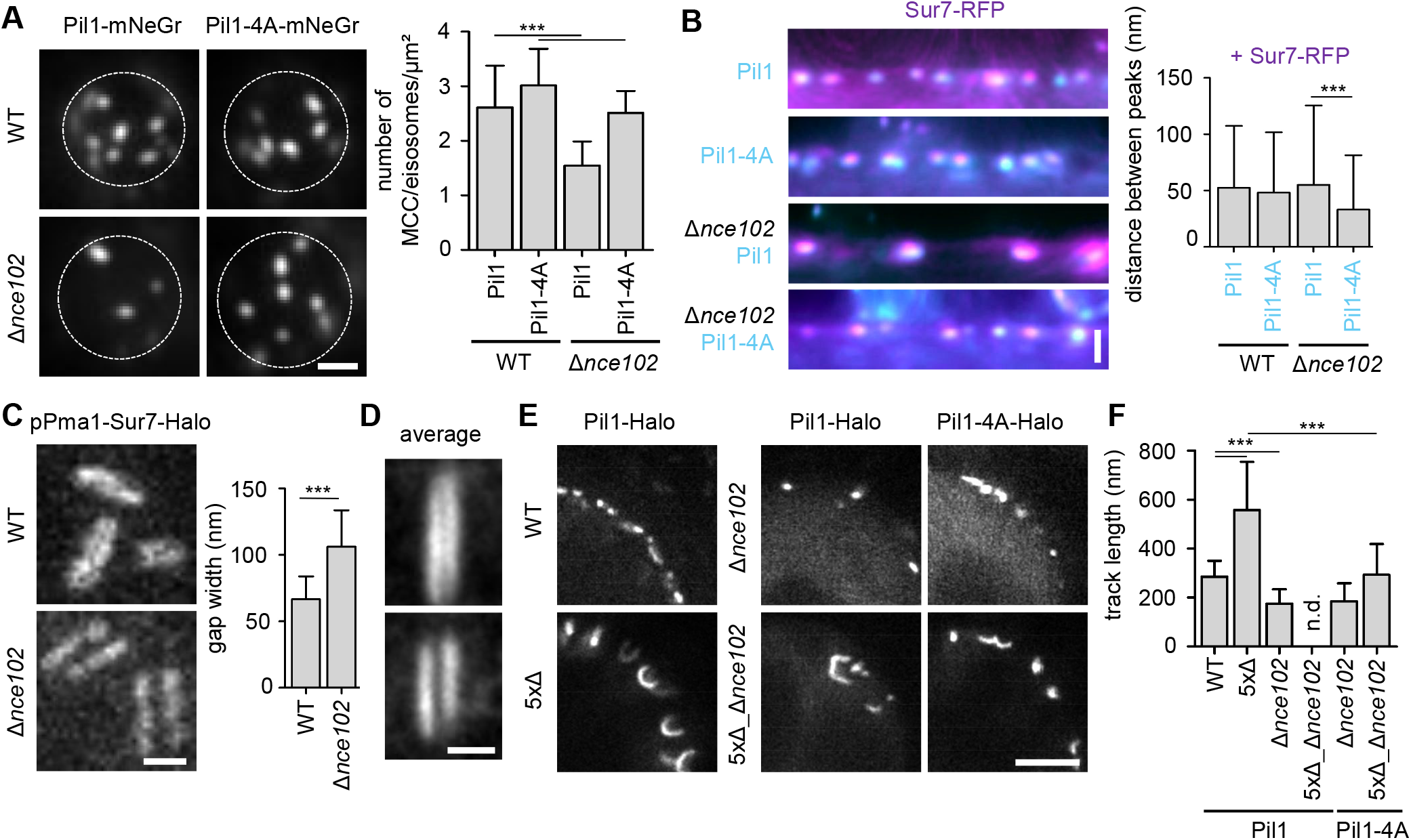
Nce102-tetraspanner contributes to the topography of MCC/eisosomes. A TIRFM images of Pil1-mNeGr or Pil1-4A-mNeGr in WT and Δ*nce102* cells. Bar graphs indicate surface density of MCC/eisosomes in number per area. B Linearized profiles of WT and Δ*nce102* cells expressing Sur7-RFP (magenta) and Pil1/Pil1-4A-mNeGr (cyan). Bar graphs indicate distances between radial profile peaks for Pil1 and Sur7. C STED images of WT and Δ*nce102* cells expressing Sur7-Halo from the Pma1 promoter. The bar graphs indicate the width of the gap between opposing Sur7-strands. D Averaged images of MCC/eisosomes marked by Sur7-Halo expressed from the Pma1 promoter in WT and Δ*nce102* cells (n = 12). E STED images of indicated mutants expressing Pil1/Pil1-4A-Halo. F Bar graphs indicate track-length of Pil1/Pil1-4A-Halo labelled structures in various mutants. Data information: (A-C) Bar graphs, unpaired t-tests, n = 42-64 cells (A), n = 92-193 MCC/eisosomes (B), n = 74-111 MCC/eisosomes (C), (F) Bar graph, ANOVA with Tukey’s multiple comparison test, n > 100 tracks. Scale bars: 1 μm, 200 nm (C, D).

### A link between lipid composition and MCC/eisosome organization

The results described above identify important roles for members of the Sur7 and Nce102 families of tetraspanners in the formation and topography of MCC/eisosomes. Generally, membrane micro- and nano-domain formation is often driven by a combination of protein-protein and protein-lipid interactions. We therefore also investigated the respective effects of various lipid classes on the formation of Sur7 strands. Removal of phosphatidylethanolamine (PE, Δ*psd1/2*) or phosphatidylserine (PS, Δ*cho1*) did not change the distribution of Sur7, as visualized either by TIRF (Fig 6A, B) or STED microscopy (Fig 6C). Depletion of phosphatidylcholine (PC, Δ*cho2Δopi3*) or acute perturbation of complex sphingolipids by treatment with AureobasidinA (AbA, Fig EV4A) led to a minor reduction of Sur7 accumulation within MCC/eisosomes (Fig 6B) but did not affect strand formation per se (Fig 6C).

**Figure 6.**
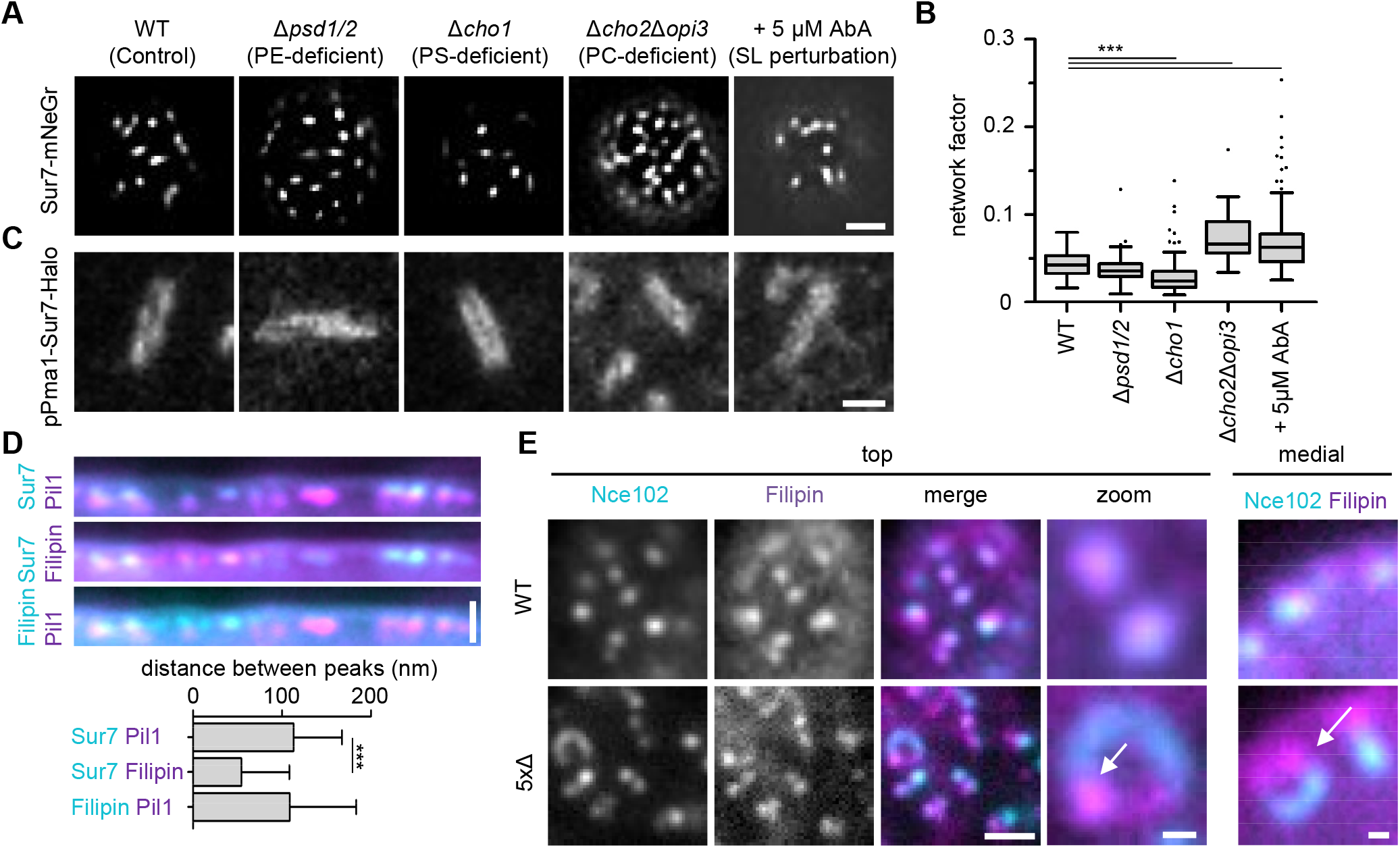
The influence of PM lipid composition on MCC/eisosome topography. A Effect of indicated lipid perturbations on lateral distribution of Sur7-mNeGr. B Lateral distribution of Sur7-mNeGr quantified by the network factor in strains shown in (A). C STED images of MCC/eisosomes marked by Sur7-Halo expressed from the Pma1 promoter in strain backgrounds shown in (A). D Linearized profiles of WT cells expressing Sur7-GFP from the Pma1 promoter (cyan) and Pil1-RFP (magenta). Strains were stained with 5 μg/ml filipin (magenta/cyan). Bar graphs indicate distances between radial profile peaks. E TIRFM images of filipin stained (magenta) WT and 5xΔ cells expressing Nce102-mNeGr (cyan). Arrows indicate filipin accumulation at ends of tubular invagination in 5xΔ cells. Data information: (B) Boxplot, ANOVA with Dunnett’s multiple comparison test, n = 30-227 cells. (D) Bar graph, ANOVA with Dunnett’s multiple comparison test, n =132-147 MCC/eisosomes. Scale bars: 1 μm, 200 nm (C, E zoom).

Previous reports have described an accumulation of sterols at MCC/eisosomes (Grossmann *et al*, 2007). We therefore tested the exact position of the ergosterol marker filipin and found that it selectively accumulates at the outer periphery of MCC/eisosomes together with Sur7 (Fig 6D). Strikingly, upon deletion of Sur7 tetraspanners filipin did not enter the closed tubes of the 5xΔ cells but remained concentrated at the ends or in the region between the ends (Fig 6E).

Our results indicate that Sur7 strand formation is not critically regulated by the presence of PE, PS and PC phospholipids or by complex sphingolipids. In addition, filipin staining indicates that regions of positive curvature at the upper edge of either MCC/eisosome furrows or tubular invaginations are enriched in ergosterol.

### The role of PIP2 in PM organization

One specific type of lipid that has previously been implicated in MCC/eisosome biogenesis is PIP2 (Fröhlich *et al*, 2014). Since this lipid is essential for yeast cells, we first asked how increased PIP2 levels affect subdomain organization of MCC/eisosomes. In agreement with previous reports (Stefan *et al*, 2002; Karotki *et al*, 2011), we found that deletion of the major PIP2 phosphatases Inp51 and Inp52 induced the formation of aberrant PM invaginations. Using the improved resolution provided by STED microscopy, we found that these structures closely corresponded to the tubes we observed in the 5xΔ strain (Fig 7A). Similar tubes were formed when increasing PIP2 levels by overexpressing the yeast PI(4)P-5 kinase Mss4 (Fig 7A). Strikingly, increased PIP2 levels and deletion of all five Sur7 family members exhibited synergistic effects on tube length (Fig 7A, B). These results suggest a close link between cellular PIP2 levels and PM domain topography driven by Sur7 tetraspanners. We therefore tested, whether PIP2 had a direct effect on localization of Sur7, as representative member of the family. Using TIRFM we found that Sur7 was in fact displaced from Nce102-labeled MCC/eisosomes under conditions in which PIP2 levels were increased (Fig 7C). More specifically, Sur7 parallel strands imaged by STED microscopy were disrupted and the tetraspanners were mislocalized to variably-shaped linear structures within the yeast PM upon deletion of *INP51* and *INP52*, irrespective of Sur7 expression levels (Fig 7D).

**Figure 7.**
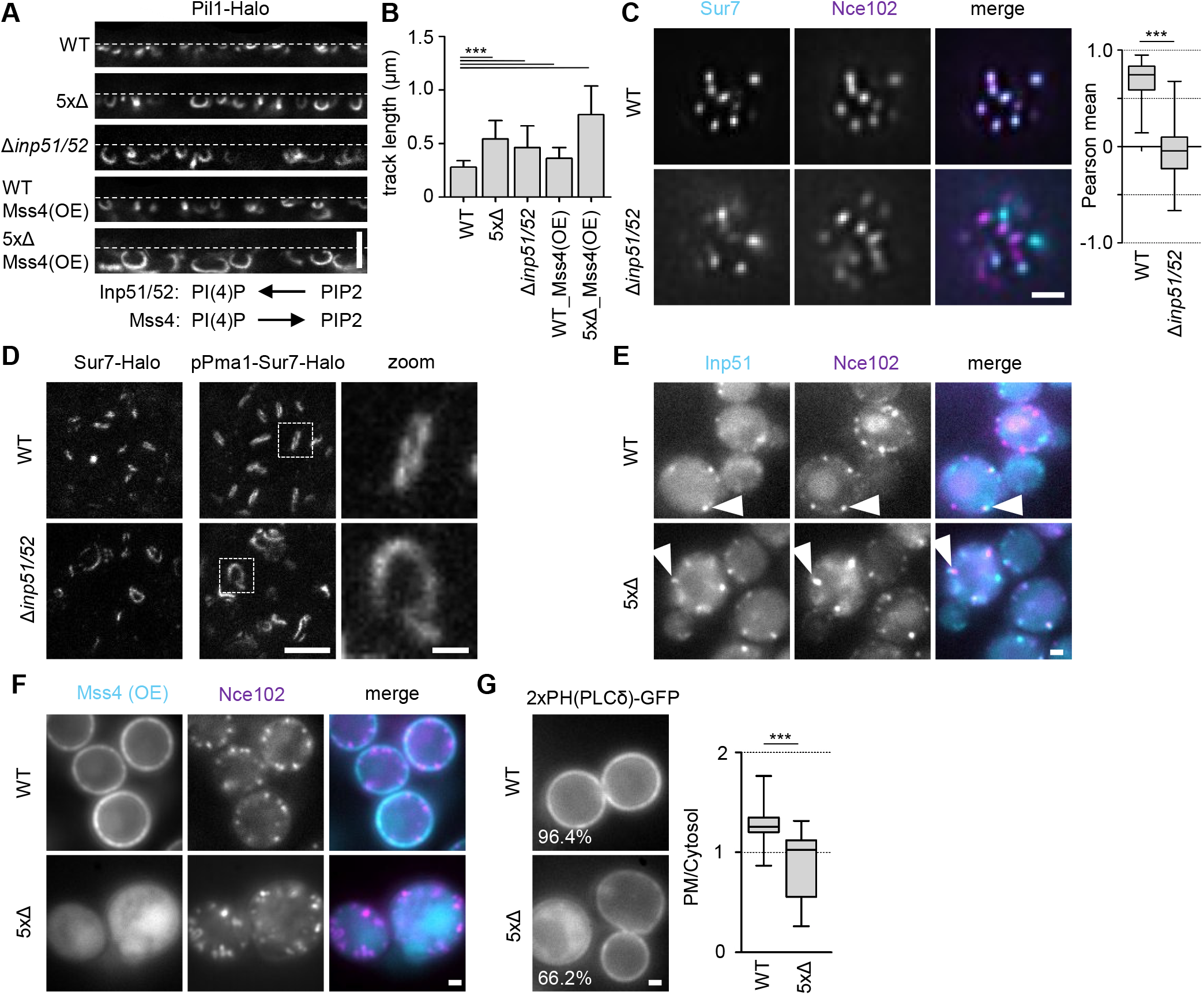
The role of PIP2 in the regulation of MCC/eisosome topography. A Linearized STED profiles of indicated mutants expressing Pil1-Halo. Mss4 was constitutively overexpressed from the GPD promoter. Dotted lines indicate PM. Function of indicated enzymes in PI(4)P- and PIP2-biosythesis is displayed schematically. B Bar graphs indicate track lengths of tubular invaginations formed under the conditions shown in (A). C Colocalization of Sur7-mNeGr (cyan) and Nce102-RFP (magenta) in MCC/eisosomes from WT and Δ*inp51/52* cells. Bar graphs indicate the respective Pearson mean coefficient. D STED images of Sur7-Halo expressed endogenously or from the Pma1 promoter in WT and Δ*inp51/52* cells. Zoomed images show details of Sur7 strand organization. E Colocalization of Inp51-mNeGr (cyan) and Nce102-RFP (magenta) in MCC/eisosomes (arrowheads) of WT and 5xΔ cells. F Localization of GFP-Mss4 overexpressed (OE) from the GPD promoter in WT and 5xΔ cells expressing Nce102-RFP (magenta). G Distribution of the PIP2 reporter 2xPH(PLCδ)-GFP in WT and 5xΔ cells. Percentage of cells that showed PM staining is indicated. Box plots show PM to cytosol ratios of GFP signal. Data information: (B) Bar graph, ANOVA with Dunnett’s multiple comparison test, n = 139-292 tracks. (C) Boxplot, unpaired t-test, n = 82-93 cells. (G) Boxplot, unpaired t-test, n > 62 cells. Scale bars: 1 μm and 200 nm (zoom)

The other known link between MCC/eisosomes and PIP2 homeostasis involves the interaction of the major PIP2 phosphatase Inp51 with Pil1 (Fröhlich *et al*, 2014). We found that the association of Inp51 with PM invaginations was maintained in 5xΔ cells (Fig 7E). The product of Inp51 activity, PI(4)P, is the substrate for Mss4 and is at the same time required for recruitment of the kinase to the PM (Ling *et al*, 2012). Interestingly, we found that overexpressed Mss4 was no longer recruited to the PM upon deletion of all Sur7 family genes (Fig 7F). This suggests a possible disruption of PI(4)P flux in the PM of 5xΔ cells. Consistent with this idea, we found that PIP2 levels in the yeast PM, marked by 2xPH(PLCδ)-GFP, were significantly reduced in 5xΔ cells (Fig 7G). Importantly, the overall cellular lipidome of the 5xΔ strain was not markedly altered compared to control cells (Fig EV4B), but for technical reasons this analysis did not include any of the phosphorylated phosphoinositide lipids.

Our results indicate close links between PIP2 homeostasis within the PM, Pil1-mediated bending of MCC/eisosomes and Sur7-mediated inhibition of membrane tubulation.

### Acute perturbation of PM organization

Our results indicate that elevated PIP2 levels in the yeast PM could induce tubulation of MCC/eisosome furrows by displacing Sur7. We wanted to test our interpretation by using acute changes in MCC/eisosome composition. A previous report showed that PIP2 clusters in the yeast PM could be rapidly induced by reducing overall PM tension with palmitoylcarnitine (PalmC, (Riggi *et al*, 2018)). We also observed the formation of such PIP2 clusters within a few minutes of exposure to 10 μM PalmC (Fig 8A) and found that these clusters often were closely associated with MCC/eisosomes marked by Nce102 (Fig 8B). We next tested the effect of PalmC treatment on Sur7 localization (again using Sur7 as representative marker for the Sur7 family). We found that after 10 min of treatment, Sur7 was still associated with Pil1-marked MCC/eisosomes, but was laterally displaced (Fig 8C). In addition, the radial distance between Pil1 and Sur7 was significantly increased, while Lsp1 and Pil1 remained closely associated (Fig 8D), indicative of deeper furrows or tubulation. Using STED microscopy we found that Sur7 strands remained intact upon PalmC-treatment but that lateral displacement was characterized by strong curvature of Sur7 strands and a frequent appearance of circular or ovoid structures (Fig 8E). Importantly, Pil1 labeled structures in PalmC-treated cells exhibited increased curvature and elongation, consistent with formation of membrane tubes. Two color imaging of Sur7 strands (STED) and Pil1 tubes (confocal) clearly confirmed the lateral segregation of the two MCC/eisosomal subdomains (Fig 8F). Displacement of Sur7 was also observed when PM tension was reduced by exposure of cells to hyperosmotic conditions (1 M sorbitol for 10 min; Fig 8G), indicating that the effect was not specific for PalmC-treatment.

**Figure 8.**
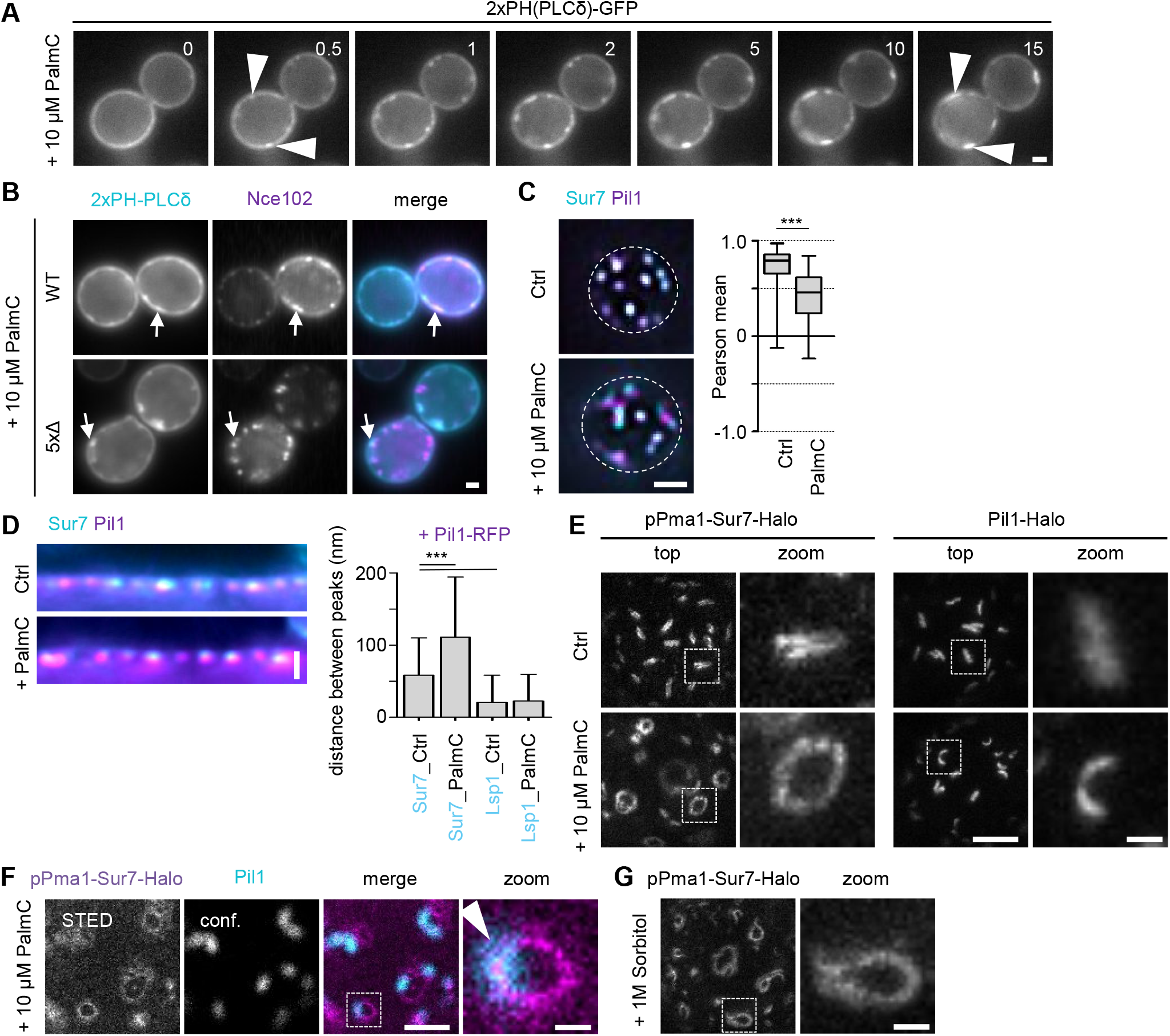
Effects of acute perturbation of MCC/eisosome topography. A Effect of addition of 10 μM palmitoylcarnitine (PalmC) on PM distribution of PIP2, marked by 2xPH(PLCδ)-GFP. Time in min after PalmC treatment. Arrowheads indicate PIP2 clusters. B Colocalization of Nce102-RFP (magenta) with PIP2 marked by 2xPH(PLCδ)-GFP (cyan) in WT and 5xΔ cells, 10 min after addition of PalmC. Arrows indicate colocalization in clusters. C Colocalization of Sur7-mNeGr (cyan) and Pil1-RFP (magenta) in WT cells before and after treatment with PalmC. Box plots indicate Pearson correlation coefficients in respective conditions. D Linearized profiles and distances of radial profile peaks between Pil1-RFP (magenta) and Sur7-mNeGr (cyan) before and after PalmC treatment. The distance between Lsp1-mNeGr and Pil1-RFP was used as control (images not shown). E STED images of WT cells expressing Sur7-Halo from the Pma1 promoter or endogenous Pil1-Halo before and after PalmC treatment. Zoom shows curved structures observed under PalmC treatment. F Colocalization of WT cell expressing Sur7-Halo from the Pma1 promoter (magenta, STED) and Pil1-mNnGr (cyan, confocal) after 10 μM PalmC treatment. Arrow head indicates the degree of overlap. G STED images of WT cells expressing Sur7-Halo from the Pma1 promoter after addition of 1 M sorbitol for 5 min. Data information: (C) Box plot, unpaired t-test, n = 37-53 cells. (D) Bar graph, ANOVA with Dunnett’s multiple comparison test, n = 112-159 MCC/eisosomes. Scale bars: 1 μm, 200 nm (zoom)

In summary, we could show that an acute reduction in PM tension and concomitantly altered PIP2 distribution leads to displacement of Sur7 from MCC/eisosomes and to a simultaneous increase of MCC/eisosome curvature and possibly tubulation.

### Mechanistic basis of Sur7 function in subdomain organization

Our results with Δ*inp51/inp52* cells suggested that correct accumulation of Sur7 in MCC/eisosomes is essential for local membrane topography. We therefore wanted to test whether artificial tethering of Sur7 to MCC/eisosomes could overcome the phenotypic changes induced by PIP2 overproduction. To this end, we fused Sur7 directly to the only membrane-resident component that we could identify inside the MCC/eisosome furrow, Nce102. Importantly, the Nce102-Sur7 chimera was functional as it rescued the tubulation phenotype of 5xΔ cells (Fig 9A, B). Remarkably, the membrane topography of Δ*inp51/inp52* cells was also completely restored (Fig 9A, B), as was the segregation of endogenous Sur7 from MCC/eisosomes (Fig 9C).

**Figure 9.**
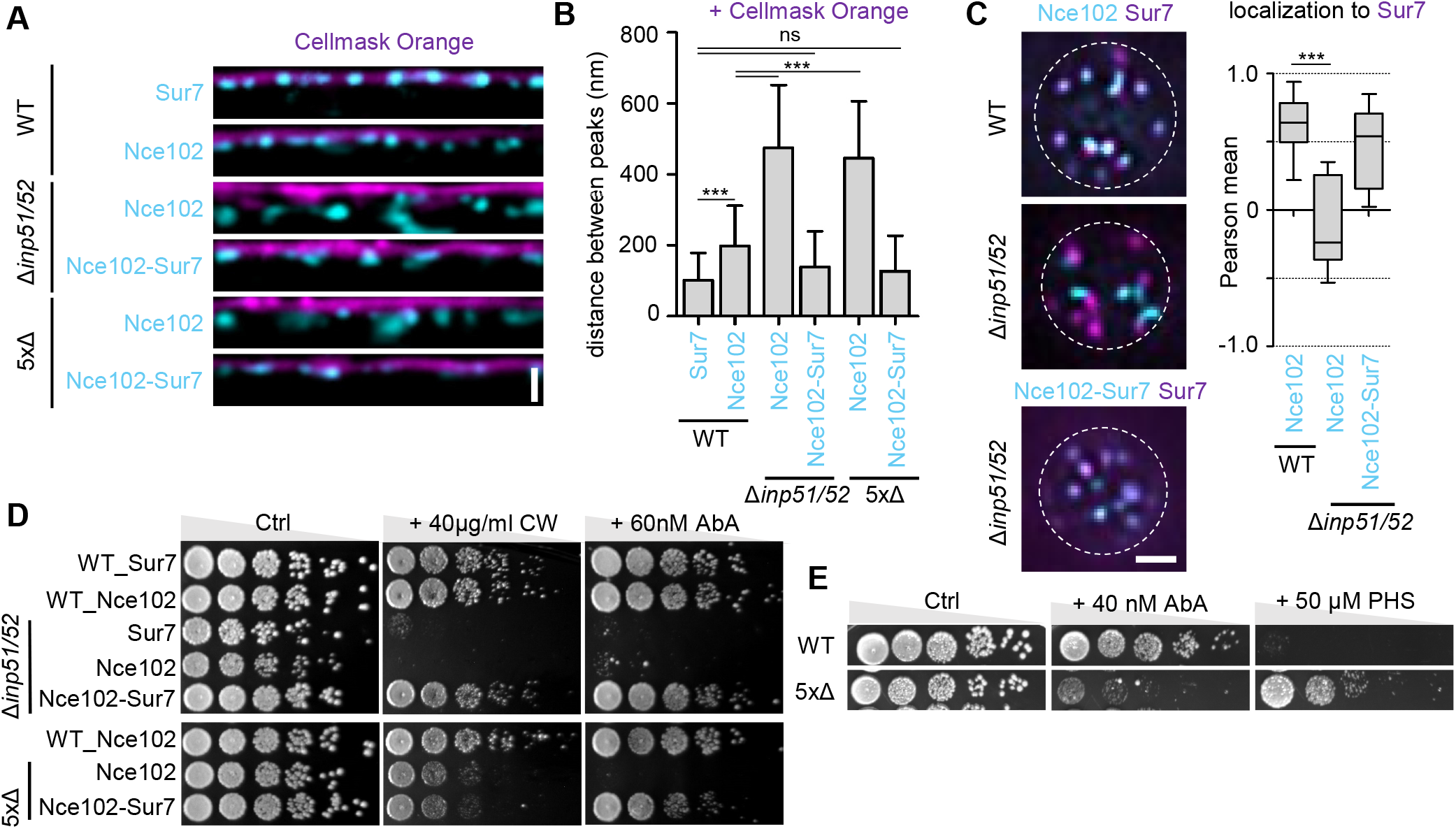
Synthetic recruitment of Sur7 controls the topography and function of MCC/eisosomes. A Linearized profiles of WT, Δ*inp51/52* and 5xΔ cells expressing mNeGr fused to Sur7, Nce102 or an Nce102-Sur7-chimera (cyan). PM labeled with Cellmask Orange (magenta). B Bar graph of distances between radial profile peaks of Cellmask Orange and proteins in (A). C Colocalization of mNeGr fusions to Nce102 or the Nce102-Sur7-chimera (cyan) with Sur7-RFP (magenta) in WT and Δ*inp51/52* cells. Quantification via Pearson correlation coefficient. D Growth assay for WT, Δ*inp51/52* or 5xΔ cells expressing mNeGr fusions to Sur7, Nce102 or the Nce102-Sur7-chimera. Five-fold serial dilutions were cultured on YPD plates containing either 40 μg/ml Calcofluor White (CW) or 60 nM AureobasidinA (AbA). E Growth assay of WT and 5xΔ cells on YPD plates containing either 40 nM AbA or 50 μM phytosphingosine (PHS). Data information: (B) Bar graph, ANOVA with Tukey’s multiple comparison test, n = 126-178 MCC/eisosomes. (C) Box plot, ANOVA with Dunnett’s multiple comparison test, n = 14-28 cells. Scale bars: 1 μm.

To explore the broader physiological consequences of altered yeast PM topography, we monitored growth of various yeast strains forming tubular invaginations in the PM. We found increased sensitivity of Inp51/52-deficient strains to cell wall stress induced by calcofluor white (Fig 9D). In addition, both Δ*inp51/52* and 5xΔ cells were hypersensitive to AbA, an inhibitor of complex sphingolipid synthesis (Fig 9D). Remarkably, hypersensitivities in either strain could be fully rescued by expression of the Nce102-Sur7 chimera in either 5xΔ or Δ*inp51/inp52* cells (Fig 9D). A potential link between MCC/eisosome topography and sphingolipid homeostasis was further underscored by a reciprocal resistance of 5xΔ cells to phytosphingosine (PHS, Fig 9E).

Taken together, we demonstrated that artificial anchoring of Sur7 to MCC/eisosomes is able to maintain normal PM topography and stress resistance in cells with increased PIP2 levels. This suggests a central function for Sur7 in regulating MCC/eisosome topography and function.

## Discussion

In this study we have uncovered opposing functions for two yeast tetraspanner families in modulating membrane curvature and domain topography at the yeast PM. More specifically, we found that the maintenance of stable PM furrows requires a balance between two force generating complexes: Negative curvature at the base of MCC/eisosome furrows is generated by PIP2 and BAR domain proteins in cooperation with the tetraspanner Nce102. This force is countered by Sur7 family tetraspanners that assemble into multimeric strands at the upper edges of furrows and prevent the transformation of PM furrows into closed membrane tubes.

Using super resolution STED microscopy, we were able to localize the different components of MCC/eisosomes with great precision. We found that Sur7 proteins form parallel strands along the longitudinal axis at the upper edges of MCC/eisosome furrows (schematic representation in Fig 10A). We observed physical interactions between various members of the Sur7-family, suggesting that the strands correspond to polymeric protein complexes. Neither the N- nor C-terminal cytosolic extensions of Sur7 were necessary for these interactions, indicating that strands were formed via transmembrane regions or extracellular loops in Sur7. Interestingly, similar polymeric structures have been described for other tetraspanner families such as the tetraspanins or claudin/occludins found in mammalian cells (Charrin *et al*, 2014; Lal-Nag & Morin, 2009). While we were able to demonstrate that the Mup1 methionine permease and the sterol dye filipin were spatially associated with the Sur7 strands, we could see a clear separation of Sur7 at the upper edge of the MCC/eisosome furrows from the basal region, which is in turn defined by the presence of Nce102 tetraspanners and the cytosolic BAR domain proteins Pil1 and Lsp1 (Fig 10A). This observation of distinct subdomains within MCC/eisosomes is consistent with previous measurements using PALM (Appadurai *et al*, 2020) and immunogold labeling (Strádalová *et al*, 2009). Importantly, Nce102 is so far the only integral PM protein that has been assigned to the basal portions of MCC/eisosome furrows.

**Figure 10.**
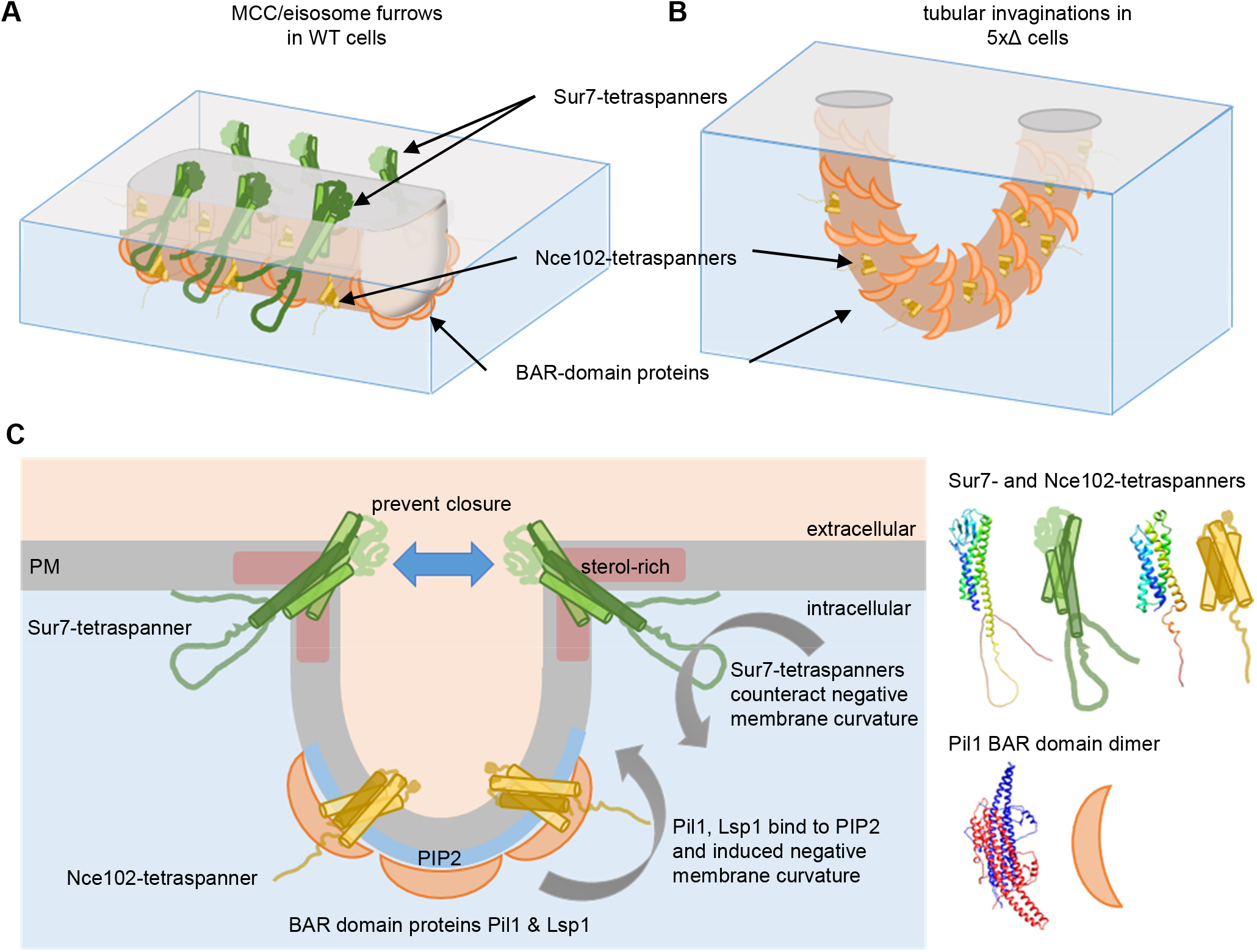
Conceptual model for tetraspanner functions in MCC/eisosome organization. A Illustration of Sur7 parallel strands at the upper edges of MCC/eisosome furrows in WT cells. B Illustration of PM topography and tubular invaginations in 5xΔ cells. C Mechanistic model depicting the balance of forces between different MCC eisosome subdomains. Negative curvature is induced by the binding of Pil1 and Lsp1 to PIP2-enriched PM domains. Positive curvature at the ergosterol-enriched upper edge of the formed furrows is stabilized by Sur7 parallel strands. Nce102 is concentrated in the lower portion of the MCC/eisosome furrow, and contributes to domain stability and shape. Grey arrows indicate direction of forces. Blue arrow illustrates the inhibition of furrow closure. Schematic protein structures are based on AlphaFold predictions.

A key result of our study is the observation that, in the absence of Sur7 tetraspanners, the typical halfpipe-shaped MCC/eisosome furrows are converted into closed membrane tubes (Fig 10B). Tube formation is dependent on the presence of the BAR domain protein Pil1. In addition, tubulation of artificial membranes was previously shown to occur by default with purified Pil1 (Karotki *et al*, 2011) or Lsp1 (Zhao *et al*, 2013). These observations suggest that multimeric strands made of multiple Sur7-family proteins counteract the bending force generated by the BAR domain coat formed by Pil1 and Lsp1 (Fig 10C, (Karotki *et al*, 2011)) *in vivo*. Our data also indicate that tubes can develop directly from regular furrows, as the end-end distance of tubes corresponded to the typical length of MCC/eisosomes. In addition, we could observe tubulation upon acute perturbation of Sur7 localization induced by PalmC treatment. These results imply that Sur7 tetraspanners play a major role during MCC/eisosome biogenesis. Interestingly, our filipin labeling results indicated that sterols specifically accumulate at the upper edges of furrows and tubes (Fig 10C), suggesting that these regions are defined by a distinct local lipid composition that could also influence tetraspanner localization.

We also found that the function of Sur7 in regulating membrane topography critically depended on its cytosolic C-terminal extension: Despite its correct localization at MCC/eisosomes and unaltered self-association in Co-IP experiments, the C-terminal truncation of Sur7 failed to prevent the formation of membrane tubes. Interestingly, the C-terminal tails of tetraspanners are known to be important sites of interaction with partner proteins and thereby promote the biological functions of claudins (Lal-Nag & Morin, 2009). Similarly, the C-terminal region of Sur7 has been shown to play a key functional role in *C. albicans* morphogenesis under various stress conditions (Lanze *et al*, 2021). Further experiments will be required to identify the relevant protein and/or lipid interaction partners of the C-terminal segment that mediate its function in regulating membrane topography.

While Sur7 tetraspanners counteract the curvature induced by Pil1, we found that Nce102 positively contributed to the formation of PM furrows and tubes. In the absence of Nce102, Sur7 strands along furrows became more separated and PM tubes formed in the absence of all 6 tetraspanners (5xΔ_Δ*nce102*) were much shorter than those found in 5xΔ cells. A key role for Nce102 in regulating MCC/eisosome topography has been previously suggested. Under glucose starvation conditions, furrows become deeper (Appadurai *et al*, 2020) and simultaneously Nce102 is upregulated (Bradley *et al*, 2009; Zahumenský *et al*, 2022). On the other hand, lipid stress such as inhibition of sphingolipid synthesis leads to redistribution of Nce102 away from MCC/eisosomes and to flattening of furrows (Fröhlich *et al*, 2009; Strádalová *et al*, 2009; Loibl *et al*, 2010). While it has been suggested that Nce102 primarily regulates the phosphorylation of Pil1, its role in membrane shaping seems to be maintained even in the presence of non-phosphorylatable variants of Pil1 (Walther *et al*, 2007) (Fig 5F). Our results therefore indicate that Nce102 may also play a structural role in MCC/eisosomes maintenance. How this might be connected to our identification of Nce102 as first and so far only integral PM protein at the basal region of MCC/eisosomes furrows, remains to be addressed.

MCC/eisosomes have been previously linked to lipid homeostasis, including regulation of PIP2 levels through Pil1-associated PIP2 phosphatase Inp51 (Fröhlich *et al*, 2014) and the control of sphingolipid metabolism via the MCC-associated sensors Slm1/2 (Berchtold *et al*, 2012) and Nce102 (Fröhlich *et al*, 2009; Zahumenský *et al*, 2022). Interestingly, previous studies have indicated that an increase in PIP2 levels leads to the formation of tubular invaginations that strongly resemble those found in 5xΔ cells (Stefan *et al*, 2002; Karotki *et al*, 2011). We have now shown that these effects on MCC/eisosome topography are accompanied by a mislocalization of Sur7. This altered distribution occurs in the Δ*inp51/52* mutant, in which PIP2 levels are increased, and could surprisingly be rescued by artificially tethering Sur7 to MCC/eisosomes via Nce102. In addition, altered PIP2 distribution induced by treatment with PalmC (Riggi *et al*, 2018) led to rapid relocation of Sur7. These experiments suggest that Sur7 strand localization and/or assembly is sensitive to the local lipid composition, in particular to PIP2 distribution. However, we cannot currently distinguish between a direct interaction of Sur7 with PIP2 and indirect consequences of PIP2 and Pil1 mediated PM curvature on Sur7 localization. Importantly, the rescue of the Δ*inp51/52* mutant was demonstrated not only for morphological changes in the PM (tubulation), but also for strain sensitivity to drugs that disrupt the cell wall or inhibit sphingolipid synthesis, indicating a restoration of physiological functions.

In summary, we have identified a central role for two tetraspanner families in regulating membrane topography at the yeast PM. We propose that the unique furrow topography of MCC/eisosomes is maintained by a balance of forces. Negative curvature is induced by the BAR domains of Pil1 and Lsp1 that cooperate with PIP2 lipids and Nce102 tetraspanners at the basal region of PM furrows. In contrast, members of the Sur7 tetraspanner family assemble into parallel strands at the ergosterol-enriched upper edge of furrows, where they maintain the typical halfpipe shape of MCC/eisosome furrows by preventing their transformation into closed tubes (Fig 10C).

## Material and Methods

### Yeast strains and plasmids

All strains in this study were derived from the *S. cerevisiae* BY4741 (Euroscarf). Genomic tagging and deletions were performed by direct integration of PCR products as described previously (Janke *et al*, 2004). The five-fold tetraspanner mutant was created by a variant of the *delitto perfetto* marker-less approach using the counterselectable *Kl*URA3 marker (Stuckey & Storici, 2013). Counterselection was performed on SCD plates containing 1 mg/ml 5-fluoroorotic acid. After loss of the *Kl*URA3 marker the following remnant sequence was left at each integration site: CGT ACG CTG CAG GTC GAC AAC CCT TAA TAT AAC TTT ATA ATG TAT GTA TAG AAG TTA TTA GGT GAT ATC AGA TCC ACT AGT GGC CTA TGC. PCR products for direct integration were generated through overlapping PCR using standard methods. All plasmids were constructed using standard molecular biology techniques. Transformation into yeast cells was performed using the LiOAc method (Janke *et al*, 2004). All plasmid sequences were verified. PCR-derived endogenous integrations were verified via colony-PCR and additional sequencing for point mutations. All strains, plasmids, oligonucleotides and linker sequences used in this study are listed in Table S1.

### Media and growth conditions

If not otherwise indicated, all yeast strains were grown overnight in standard Yeast extract Peptone Dextrose medium (YPD) or in synthetic complete media with 2% glucose (SCD) at 30°C. Before imaging, cells were washed (1 min at 1000 x g) in H_2_O and diluted 1:20 in appropriate SCD medium, and grown to logarithmic phase for a further 2-4 h. The Δ*cho1*-mutant was grown in medium supplemented with 1 mM ethanolamine, and the Δ*psd1/2*-mutant was supplemented with 1 mM choline. For perturbation of complex SL-synthesis, the SCD medium was supplemented with 5 μM AureobasidinA (Clontech), diluted from a 5 mM stock solution (in ethanol) in SCD medium and cells were incubated for 1 h at 30°C. For acute modulation of the lipid composition of the PM, palmitoyl-DL-carnitine chloride (PalmC, Sigma-Aldrich) was added from a 10 mM stock (in DMSO) to a final concentration of 10 μM, and cells were incubated for the indicated time at 30°C. Hyperosmotic shock was applied by addition of 1M sorbitol for 5-10 min before imaging.

### Plate growth assays

Indicated yeast strains were grown overnight in YPD at 30°C. Cells were washed (1 min at 1000 x g) once in H_2_O and diluted 1/10 into fresh medium for further incubation over 2-3h at 30°C. Logarithmically growing cells were spotted in a five-fold serial dilution, starting at OD_600_ of 0.05, on agar plates containing indicated concentration of AureobasidinA (Clontech), Calcofluor White (Sigma-Aldrich) or phytosphingosine (ChemCruz).Plates were incubated at 30°C and imaged after 48 h.

### Filipin staining

1 ml aliquots of logarithmic grown yeast cells were washed (1 min at 1000 x g) once in PBS and resuspended in 1 ml PBS containing 5 μg/ml filipin (Sigma-Aldrich, Stock: 5 mg/ml in DMSO). Cells were incubated for 5 min at room temperature in the dark, washed once in PBS and imaged with 355 nm excitation.

### CellMask Orange staining

1 ml aliquots of logarithmic grown yeast cells were washed (1 min at 1000 x g) once in PBS and resuspended in 1 ml PBS containing 0.5 μg/ml CellMask™ Orange (stock 5 mg/ml in DMSO, ThermoFisher Scientific). Samples were incubated for 5 min at room temperature and subsequently washed three times in PBS. Cells were imaged at 561 nm excitation.

### Co-Immunoprecipitation (Co-IP)

Co-IP experiments were performed as described previousy (Bonifacino *et al*, 2001). In brief, 30 μl Protein G Sepharose®(#GE17-0618-01, Merck) was coupled to 1 μg monoclonal mouse anti-GFP antibody (#11814460001, Roche) and incubated with protein extracts. Crude extracts were obtained by crushing 50 OD_600_ cells with glass beads in non-denaturing lysis buffer (50 mM Tris/HCl pH 7.4, 300 mM NaCl, 10% Glycerin, 1% Triton X-100, 100 mM PMSF, 2 μg/ml leupeptin, 1x complete EDTA-free inhibitor cocktail). After centrifugation at 16.000 x g for 15 min at 4°C, the supernatant was pre-cleared with uncoupled Protein G Sepharose for 1 h at 4°C. An aliquot of the supernatant was diluted 1:1 with HU-Buffer + 1.5% DTT (0.2M Tris/HCl pH 6.8, 8 M Urea, 5% SDS, 1 mM EDTA, 0,1% bromophenol blue) and used as the input-sample (I-sample). The pre-cleared supernatant was split into two parts that were either incubated overnight at 4°C with Protein G Sepharose conjugated to unspecific-IgG (Control, C-sample) or to anti-GFP antibody (Co-IP, IP-Sample). The antibody-conjugated beads were washed three times with washing buffer (0.1% (w/v) Triton X-100, 50 mM Tris/HCl pH 7.4, 300 mM NaCl, 5 mM EDTA) on ice, diluted 1:1 with HU-buffer + 1.5% DTT, heated at 65°C for 30 min and analyzed via SDS-PAGE and Western blot. Membranes were probed with primary polyclonal rabbit anti-HA and anti-GFP antibodies (#51064-2-AP, #50430-2-AP, Proteintech) used in Fig EV1F and Fig 1 or mouse anti-GFP antibody (#11814460001, Roche). Signals were detected by chemiluminescence following incubation with AffiniPure Goat Anti-Rabbit or AffiniPure Goat Anti-Mouse IgG conjugated to horseradish peroxidase, respectively (#111-035-003, #115-035-003, Jackson Immuno Research).

### Protein extraction

1 ml of yeast cells (OD_600_ = 1) from logarithmic growth phase were harvested by centrifugation (1 min at 1000 x g). Cell pellets were suspended in 100 μl ice-cold H_2_O. Cell lysis and protein precipitation was performed by sequential addition of 50 μl of 2 M NaOH solution and 50 μl of a 50% TCA solution for 10 min on ice each. Samples were centrifuged for 5 min at 18,000 x g (4°C) and the pellet was resuspended in 100 μl HU-Buffer + 1.5%DTT. Samples were further processed by SDS-PAGE and immunoblotting. Samples were probed with monoclonal mouse anti-mNeonGreen (mNeGr) IgGs (#32F6, Chromotek) and monoclonal mouse anti-GAPDH IgGs (#ab125247, Abcam). Primary antibodies were detected and quantified by Western blotting, as described in the previous section.

### Lipidomics

The lipidomics data were acquired in cooperation with Lipotype GmbH (Dresden, Germany). In brief, samples (20 OD600) of yeast cells in logarithmic phase were lysed with glass beads in HPLC-grade H_2_O. Cell extracts were snap-frozen in liquid nitrogen and shipped on dry ice to Lipotype for sample extraction and analysis. Lipidome data were visualized and analyzed by Lipotype Zoom.

### Fluorescence microscopy

Coverslips were cleaned by sequential sonication in absolute Ethanol, Acetone, 1 M NaOH and H_2_O for 30 min each. Before imaging, coverslips were coated with 12 μl of 1 mg/ml Concanavalin A (Sigma-Aldrich) and air-dried. Epifluorescence (medial view) and Total Internal Reflection Fluorescence Microscopy (TIRFM, top view) was performed on iMIC-based microscopes (FEI/Till Photonics) equipped with Olympus 100x/1.45 NA oil immersion objectives and DPSS lasers at 488 nm (Cobolt Calypso, 75 mW) and 561 nm (Cobolt Jive, 150 mW). For filipin imaging a polychrome at 355 nm excitation was used. A two-axis galvanometer-driven scanning head was used to adjust TIRFM-angles individually for each color. Two separate dichroic filter cubes were used for detection of GFP and RFP signals. For filipin imaging a dichromatic quadband dichroic filter cube (zt405/488/561/640 RPC) was used. Images were acquired on an Andor iXON DU-897 EMCCD or IMAGO-QE camera controlled by the LiveAcquisition software (FEI/Till Photonics).

### STED super-resolution microscopy

For STED imaging proteins of interest were genetically fused to the HaloTag (Halo) and subsequently labelled with Janelia Fluor®646 HaloTag ligand (#GA1121, Promega) by incubating 200 nM ligand (from 200 μM stock in DMSO) was incubated with cells in SCD medium at 30°C for 30 −120 min. STED super-resolution imaging was performed on a STEDYCON scanner with pulsed 450 nm (confocal), 640 nm excitation lasers and a pulsed 775 nm depletion laser (Abberior Instruments GmbH, Göttingen, Germany). The STEDYCON was attached to a Nikon Eclipse Ti-E microscope with a 100x/1.45 NA oil immersion objective. Depletion laser power was set to obtain a pixel size of 25 nm, the pinhole was fixed at 1.1 Airy units and the pixel dwell time was set to 10 μs with five line accumulations. The STED signal was collected by an avalanche photo diode after passing a 675/25 nm bandpass filter with a gating of 1.0 - 7.0 ns. The STEDYCON was controlled via the STEDYCON Smart Control software. Samples in Fig EV2B, Fig 9F and the Δ*inp51/52* cells in Fig 7A were analyzed on a Leica TCS SP8 STED3x microscope equipped with a 100x/1.40 oil immersion objective and a pulsed (80 MHz) white light excitation laser. Excitation was performed using 488/561/633 nm laser lines and for depletion a pulsed laser at 775 nm was used. The depletion laser power was set to achieve a pixel size of 25 nm with eight line accumulations, the pinhole was fixed at 1.0 Airy units and the pixel dwell time was set to 8.6 μs. The STED signal was collected at 648-701 nm using a hybrid detector with a gating time of 0.6 – 6.0 ns. The microscope was controlled by LAS X software.

### Freeze fracture transmission electron microscopy

5 ml of yeast cells from logarithmic growth phase (4 h) were harvested by centrifugation (1 min at 1000 x g) and washed in KPi-buffer (50 mM potassium phosphate buffer, pH 5,5). Cells were fixed for 30 min in 1% (final v/V) glutaraldehyde and subsequently washed three times in KPi-buffer. Fixed samples were stored overnight in KPi-buffer with 20% BSA at 4°C. A 2 μl aliquot was loaded into a gold-coated copper carrier (3 mm ø) with a dimple, frozen in liquid ethane (−170°C) and transferred into liquid nitrogen. The sample was channeled into a Leica ACE900 and adjusted at −130°C. The sample was cut with the fracturing knife at −110°C and immediately coated with 2.5 nm Pt/C (45°, without rotation). The replica was stabilized with 30 nm C-coating (90°, 120 rpm). The replica was cleaned three times in _dd_H_2_O, in 48% H_2_SO_4_ for ~16 h (overnight), in 75% H_2_SO_4_ for 3 h and in _dd_H_2_O (5x). The sample was loaded onto a pioloform-coated copper grid and images were acquired on a Phillips CM 10 transmission electron microscope and a TEMCam F-416 camera from TVIPS (Gauting, Germany).

### Ultrathin section transmission electron microscopy

5 ml of yeast cells from logarithmic growth phase (4 h) were harvested by centrifugation (1 min at 1000 x g) and washed in phosphate buffer (PBS, pH 7.3). Cells were fixed in 2.5 % (v/V) glutaraldehyde in phosphate buffer for ~ 16 h at 4°C. After washing three times with Sörensen phosphate buffer (pH 7.3), yeast cells were post-fixed in Sörensen phosphate buffer (pH 7.3) containing 1% osmium tetroxide (OsO4) for 1 h at room temperature.

The samples were dehydrated at room temperature by passage through a graded ethanol series (30%, 50%, 70% for 10 min each and 90% and 2x 100% absolute ethanol for 15 min each), and further dehydrated in propyleneoxide for 20 min. Subsequently, the samples were infiltrated in a 2:1 (v/V) mixture of propyleneoxide: epon (overnight, 4°C), 1:1 mixture (3h, room temperature) and 1:2 (overnigth, 4°C). Finally, the samples were infiltrated with pure epon for 3 h and the again overnight. Infiltrated samples were polymerized in epon-filled molds at 60°C for 36 h. Ultrathin sections (~ 60 nm) were cut on an ultramicrotome (Reichert Ultracut S) with a diamond knife. Sections were placed on pioloform-coated TEM copper grids and contrasted with uranyl acetate (20 min) and lead-citrate (70-90 sec.). Images were acquired on a Phillips CM 10 transmission electron microscope with a TEMCam F-416 camera from TVIPS (Gauting, Germany).

### Image processing and visualization

TIRFM images were processed using Fiji and MATLAB. Raw TIRFM images were deconvolved (deconvlucy) using the Lucy-Richardson algorithm with 20 iterations and a PSF function obtained from 100 nm tetraspec microspheres in MATLAB. Protein colocalization (Pearson mean) and fluorescence intensity distribution (Network factor) were calculated from two-color TIRFM images using a customized script in MATLAB. In brief, cells were automatically detected, deconvolved and thresholded. For colocalization between GFP and RFP signals, a mask was generated for each channel and combined by AND-function. The colocalization between both channels was calculated using the Pearson correlation coefficient.

The normalized intensity distribution (i.e., the network factor) (Spira *et al*, 2012; Busto *et al*, 2018) was calculated from deconvolved TIRFM images. A rolling ball filter with 25 pixel diameter was used for background equalization. Number of iterations for deconvolution was set to 20. Low values (≤ 0.15) indicate clustered and patchy structures, whereas higher values (> 0.15) represent a more disperse and network-like distribution.

Equatorial epifluorescence images and STED images are shown as raw images. All images were contrast-adjusted and zoomed for presentation purposes only. Additionally, samples treated with CellMask™ Orange were denoised, using the “remove background” algorithm (radius 10 pixel) in Fiji.

Linearized profiles of single yeast cells were obtained from circular segmented ROIs along the cell periphery. The ROIs were processed with the “straightener” function in Fiji. For visualization, images were scaled with bilinear interpolation and X, Y scaling-factor of 4. Linearized profiles were used exclusively for visualization and not for quantification of intensity profiles.

The radial distance between two signal peaks was measured using a semi-automated Fiji macro. Two-color images were merged and shift-corrected. Subsequently, three linear ROIs perpendicular to individual MCC/eisosomes were drawn per mother cell at intervals of approximately 120 degrees. ROIs were further processed by using the “multichannel plot profile” function in Fiji. In empty controls (Fig 4A, C), the edge green autofluorescence signal in the cytosol was used as reference for the extracellular position in the “empty” control (Fig 4A, C). Distances were transferred and visualized in Prism 5.0 (GraphPad).

To calculate the PM/cytosol ratio, two circular ROIs were drawn around the cell periphery enclosing the PM. The PM intensity was obtained by subtracting inner from outer ROI using a custom Fiji macro. Background values were subtracted before calculating the ratios.

### Protein structure prediction

Protein topology was predicted by TMHMM (Krogh *et al*, 2001). Protein structures were obtained from the Alphafold structure database (Jumper *et al*, 2021; Varadi *et al*, 2022).

## Data information and statistics

Boxplots show min to max spread (whiskers) and indicate median values (line in boxes). Bar graphs depict mean ± standard deviation (SD). Number of measurements (n) is indicated in each figure legend. Statistical analysis was carried out with Prism 5.0 (GraphPad). For statistical comparison between two samples, an unpaired t-test (*** p < 0.001) was used. For comparison of multiple samples with a single control, one-way ANOVA with Dunnett’s multiple comparison test (*** p < 0.01) was used. For comparison of multiple columns, one-way ANOVA with Tukey’s multiple comparison test (*** p < 0.01) was performed. All graphs were generated in Prism 5.0 (GraphPad).

## Acknowledgements

We are grateful to Florian Fröhlich for helpful suggestions regarding lipid effects on PM organization. We thank Jens Wendt for help with Fiji Macros and Jon V Busto for generating several plasmids. We thank Florian Fröhlich, Paul Hardy and Volker Gerke for valuable comments on the manuscript. This work was supported by the German Research Foundation (SFB944 and SFB1348 to RW-S and JK).

## Author contributions

DH, RW-S and CS conceived the project, designed experiments, and analyzed data. DH, JW and AE conducted experiments and analyzed data. UK, CR and JK planned EM experiments. CR and UK performed electron microscopy. MK and JK supervised and helped DH carry out STED nanoscopy. AJ performed Co-IP experiments. DH and RW-S wrote the manuscript with input from all authors.

---

**Figure EV1.**
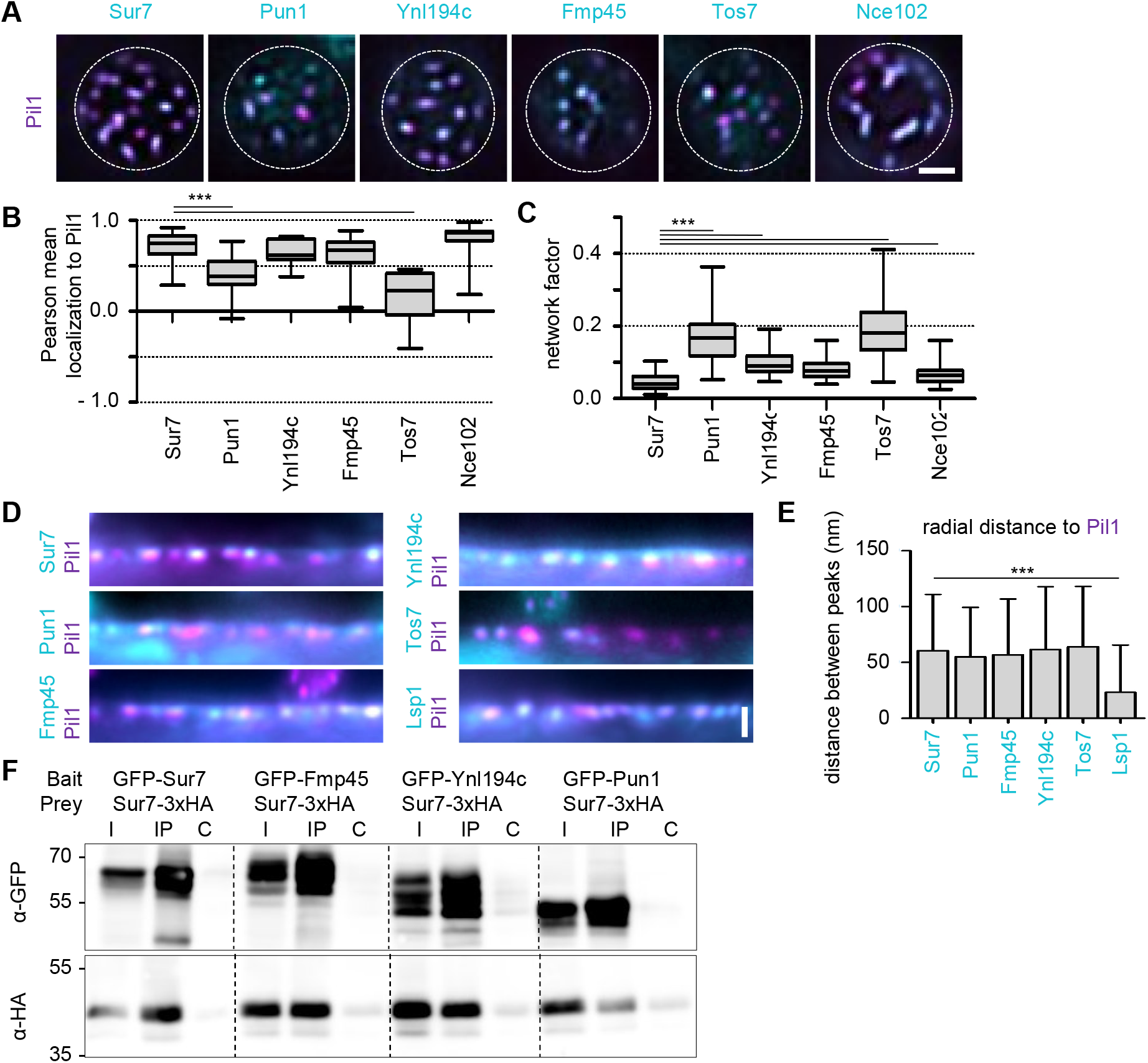
Yeast tetraspanners define subdomains in MCC/eisosomes - Supplement. A TIRFM images showing colocalization of indicated tetraspanners (C-terminally endogenously fused to GFP (cyan)) with the MCC/eisosome marker Pil1-RFP (magenta). B Pearson correlation coefficients for colocalization shown in (A). C Lateral distribution of tetraspanners from (A) quantified using the network factor. D Linearized profiles of WT cells expressing endogenous mNeGr fusions to indicated Sur7 tetraspanners and Lsp1 (cyan) together with Pil1-RFP (magenta). E Quantification of radial distances between indicated protein pairs shown in E. F Co-Immunoprecipitation of different Sur7 tetraspanner pairs. The indicated GFP-tagged bait proteins were overexpressed from the GPD promoter and were pulled down with Anti-GFP. The prey protein Sur7-3xHA was expressed under the Pma1 promoter. Western blot probed with an antibody directed against HA and GFP. I: Input, IP: Co-IP with anti-GFP, C: IP with unspecific IgG. Data information: (B, C) Boxplots, ANOVA with Dunnett’s multiple comparison test, n = 29-49 cells (B), n = 59-150 cells (C). (F) Bar graph, ANOVA with Dunnett’s multiple comparison test, n = 32-72 MCC/eisosomes. Scale bars: 1 μm.

**Figure EV2.**
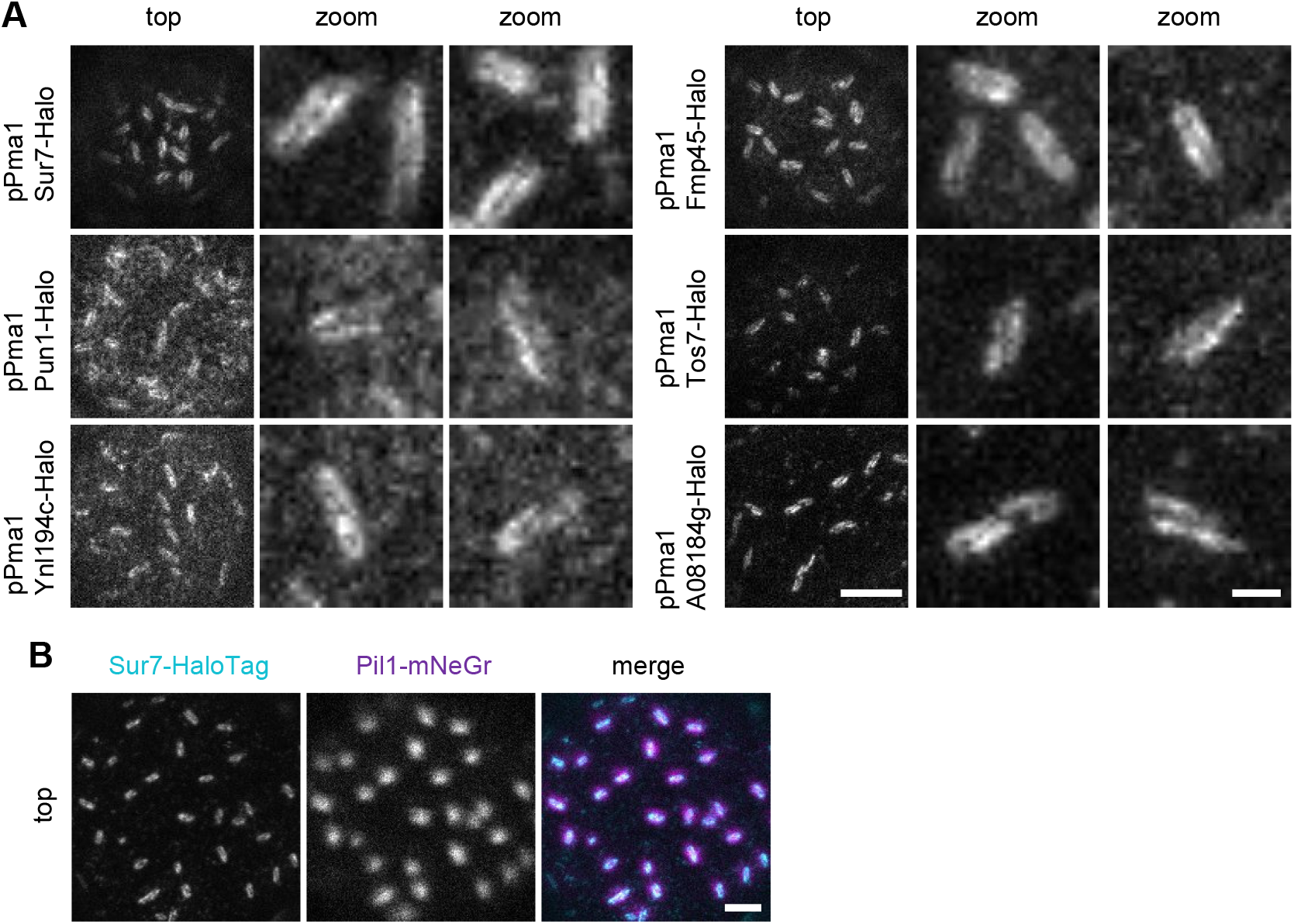
Super-resolution imaging of Sur7-tetraspanners - Supplement. A STED microscopy of indicated tetraspanners expressed from the Pma1 promoter and fused to Halo. B Two-color STED and confocal images of WT cells expressing endogenously tagged Sur7-Halo (STED, magenta) and Pil1-mNeGr (confocal, cyan). Data information: Scale bars: 1 μm (A, B), 200 nm (zoom)

**Figure EV3.**
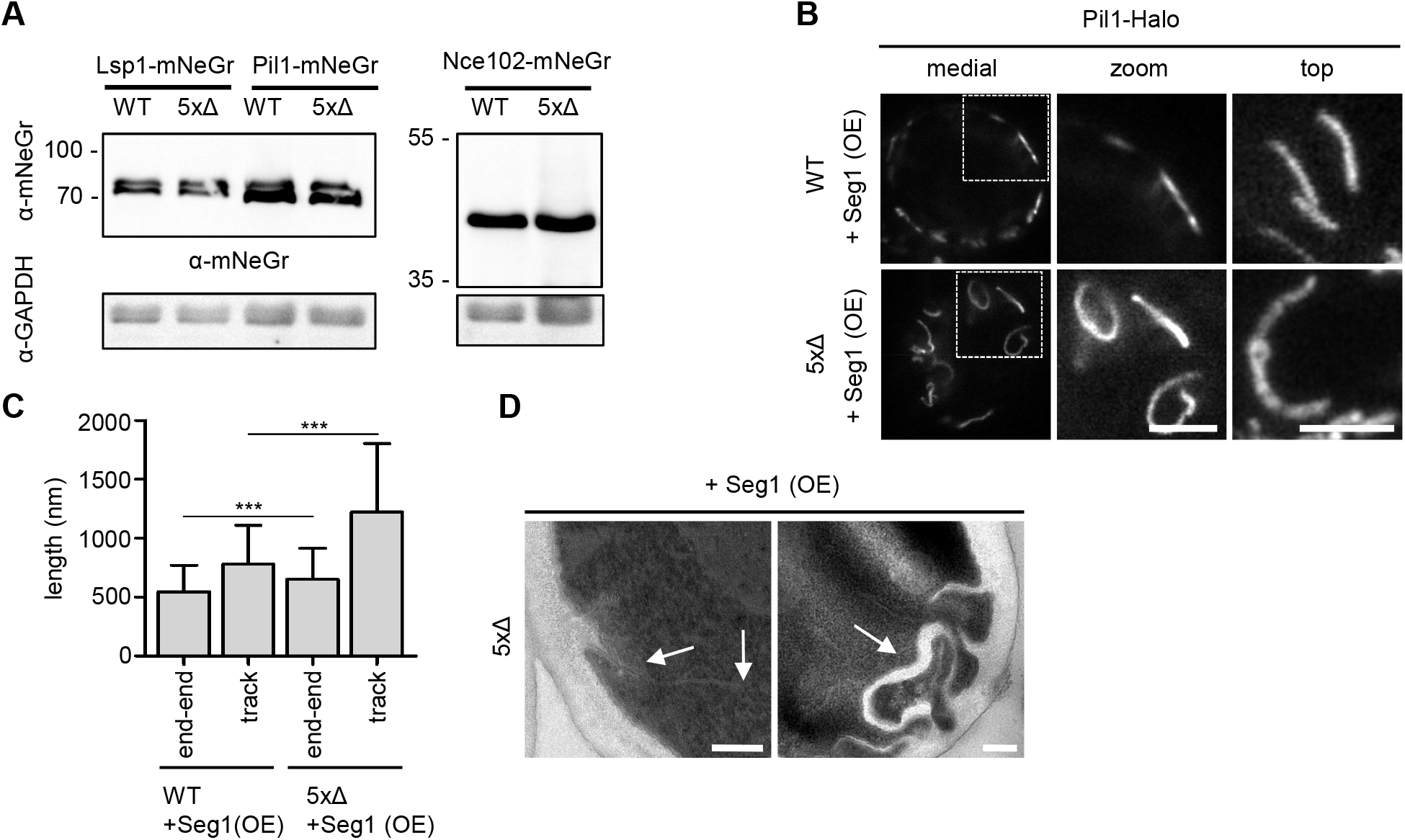
Sur7-tetraspanners prevent closure of MCC/eisosome furrows - Supplement. A Western blot analysis depicting the expression of mNeGr fusions to Lsp1, Pil1 and Nce102 in WT vs 5xΔ cells. B STED microscopy of endogenously tagged Pil1-Halo in WT and 5xΔ cells that overexpress Seg1 from the GPD promoter (OE). C Quantification of track length for Pil1-Halo positive structures in (B). D TEM micrographs of ultrathin sections of 5xΔ cells overexpressing Seg1 from the GPD promoter (OE). Arrows indicate PM invaginations. Data information: (C) Bar graph, unpaired t-test, n = 53-175 tracks. Scale bars: 1 μm, 200 nm (D)

**Figure EV4.**
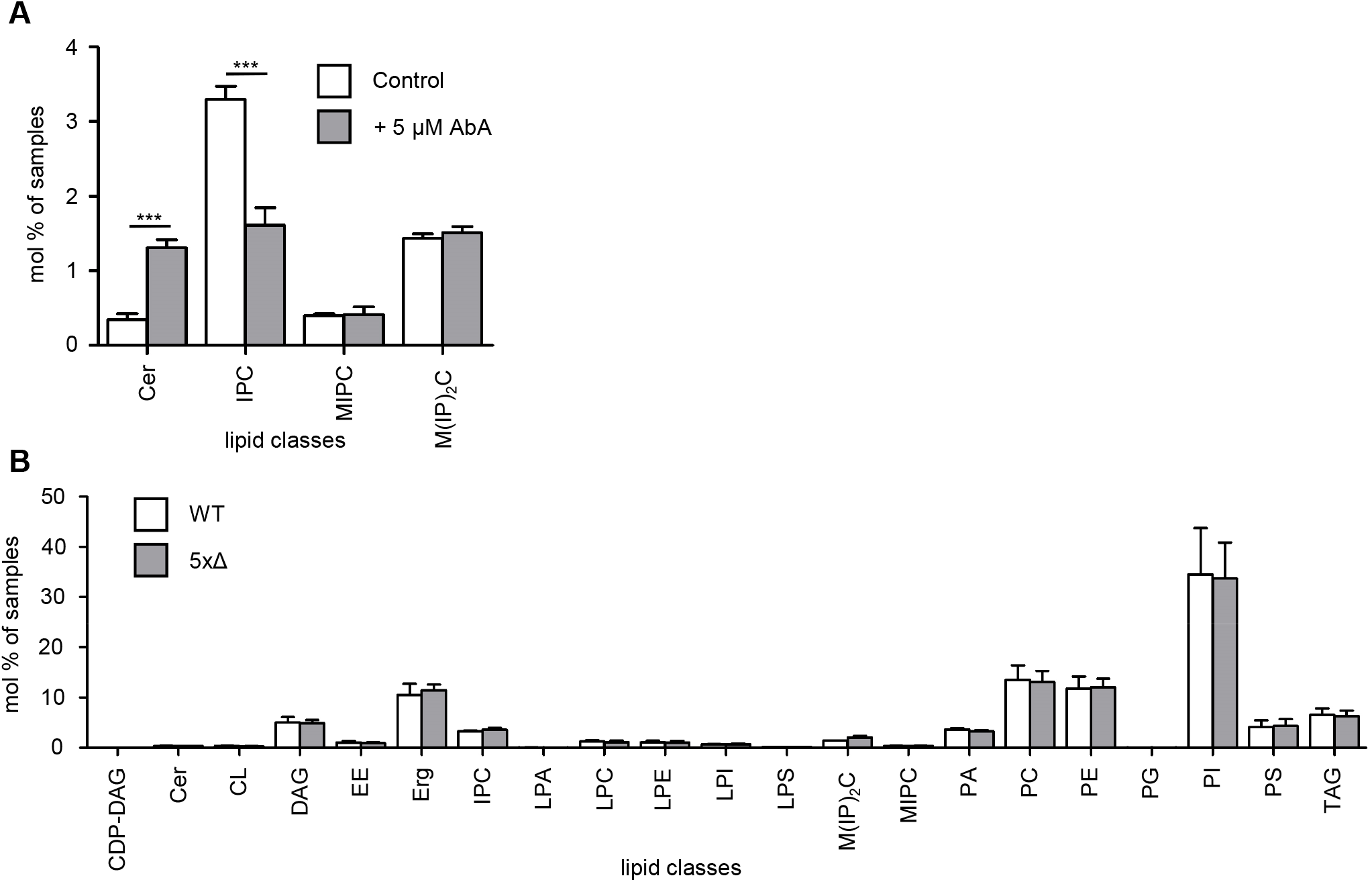
The influence of PM lipid composition on MCC/eisosome topography - Supplement. A Lipidomic analysis of sphingolipids from whole cell extracts of WT cells. Comparison of lipids identified in untreated cells vs. cells cultured in 5 μM AureobasisinA (AbA) for 1 h. B Extended lipidomic profiles for whole cell extracts of WT and 5xΔ cells. Abbreviations: CDP-DAG, cytidine diacylglycerol; Cer, ceramide; CL, cardiolipin; DAG, diacylglycerol; EE, ergosterol esters; Erg, ergosterol; IPC, Inositolphosphoryl-ceramides; LPA, lysophosphatidic acid; LPC, lysophosphatidylcholine; LPE, lysophosphatidylethanolamine; LPI, lysophosphatidylinositol; LPS, lysophosphatidylserine; M(IP)2C, mannosyl-di-(inositol-phosphoryl)-ceramides; MIPC, mannosyl-inositolphosphoryl-ceramides; PA, phosphatidic acid; PC, phosphatidylcholine; PE, phosphatidylethanolamine; PG, phosphatidylglycerol; PI, phosphatidylinositol; PS, phosphatidylserine; TAG, triacylglycerol Data information: (A) Bar graph, unpaired t-test, N = 3.

| **Internal reference** | **Strain** |
| --- | --- |
| GFP fusion library | EY0986 (BY4741 - ATC) |
| RWS2255 | BY4741 |
| RWS2256 | BY4742 |
| RWS3149 | BY4741 |
| RWS3176 | BY4741 |
| RWS3179 | BY4741 |
| RWS3185 | BY4741 |
| RWS3321 | BY4741 |
| RWS3322 | BY4741 |
| RWS3323 | BY4741 |
| RWS3340 | BY4741 |
| RWS3795 | BY4741 |
| RWS3797 | BY4741 |
| RWS3838 | BY4741 |
| RWS3949 | BY4741 |
| RWS3951 | BY4741 |
| RWS4045 | BY4741 |
| RWS4053 | BY4741 |
| RWS4170 | BY4741 |
| RWS4246 | BY4741 |
| RWS4248 | BY4741 |
| RWS4249 | BY4741 |
| RWS4250 | BY4741 |
| RWS4266 | BY4741 |
| RWS4278 | BY4741 |
| RWS4280 | BY4741 |
| RWS4285 | BY4741 |
| RWS4286 | BY4741 |
| RWS4290 | BY4741 |
| RWS4292 | BY4741 |
| RWS4304 | BY4741 |
| RWS4305 | BY4741 |
| RWS4306 | BY4741 |
| RWS4307 | BY4741 |
| RWS4308 | BY4741 |
| RWS4309 | BY4741 |
| RWS4323 | BY4741 |
| RWS4367 | BY4741 |
| RWS4369 | BY4741 |
| RWS4422 | BY4741/BY4742 |
| RWS4445 | BY4741 |
| RWS4450 | BY4741 |
| RWS4451 | BY4741 |
| RWS4453 | BY4741 |
| RWS4454 | BY4741 |
| RWS4455 | BY4741 |
| RWS4484 | BY4741 |
| RWS4507 | BY4741 |
| RWS4588 | BY4741 |
| RWS4589 | BY4741 |
| RWS4594 | BY4741 |
| RWS4595 | BY4741 |
| RWS4598 | BY4741 |
| RWS4832 | BY4741 |
| RWS4834 | BY4741 |
| RWS4872 | BY4741 |
| RWS4918 | BY4741 |
| RWS4958 | BY4741 |
| RWS4961 | BY4741 |
| RWS4970 | EY0986 (BY4741 - ATC |
| RWS4971 | EY0986 (BY4741 - ATC |
| RWS4977 | BY4741 |
| RWS4978 | BY4741 |
| RWS4999 | BY4741 |
| RWS5000 | BY4741 |
| RWS5035 | BY4741 |
| RWS5037 | BY4741 |
| RWS5038 | BY4741 |
| RWS5039 | BY4741 |
| RWS5041 | BY4741 |
| RWS5080 | BY4741 |
| RWS5087 | BY4741 |
| RWS5088 | BY4741 |
| RWS5097 | BY4741 |
| RWS5122 | BY4741 |
| RWS5131 | BY4741 |
| RWS5132 | BY4741 |
| RWS5133 | BY4741 |
| RWS5134 | BY4741 |
| RWS5168 | BY4741 |
| RWS5199 | BY4741 |
| RWS5342 | BY4741 |
| RWS5343 | BY4741 |
| RWS5344 | BY4741 |
| RWS5368 | BY4741 |
| RWS5369 | BY4741 |
| RWS5374 | BY4741 |
| RWS5399 | BY4741 |
| RWS5401 | BY4741 |
| RWS5402 | BY4741 |
| RWS5409 | BY4741 |
| RWS5410 | BY4741 |
| RWS5411 | BY4741 |
| RWS5412 | BY4741 |
| RWS5424 | BY4741 |
| RWS5432 | BY4741 |
| RWS5442 | BY4741 |
| RWS5457 | BY4741 |
| RWS5459 | BY4741 |
| RWS5460 | BY4741 |
| RWS5461 | BY4741 |
| RWS5473 | BY4741 |
| RWS5488 | BY4741 |
| RWS5545 | BY4741 |
| RWS5547 | BY4741 |
| RWS5551 | BY4741 |
| RWS5557 | BY4741 |
| RWS5560 | BY4741 |
| RWS5561 | BY4741 |
| RWS5604 | BY4741 |
| RWS5612 | BY4741 |
| RWS5613 | BY4741 |
| RWS5625 | BY4741 |
| RWS5636 | BY4741 |
| RWS5661 | BY4741 |
| RWS5663 | BY4741 |
| RWS5665 | BY4741 |
| RWS5675 | BY4741 |
| RWS5679 | BY4741 |
| RWS5681 | BY4741 |

## Genotype/description

MATa his3Δ1 leu2Δ0 met15Δ0 ura3Δ0

BY4741: MATa; His3Δ 1; Leu2Δ 0; met15Δ 0; Ura3Δ 0

BY4742: MATα; His3Δ 1; Leu2Δ 0; lys2Δ 0; Ura3Δ 0

Sur7-GFP::HIS3MX6_pTL58-pPma1-Sur7-3xHA::Leu2

Pil1-GFP::HIS3MX6_pTL58-pPma1-Sur7-3xHA::Leu2

pTL58-pPma1-Sur7-3xHA::Leu2

Mup1ΔC-GFP::hphNT1_pTL58-pPma1-Sur7-3xHA::Leu2

Pil1-mRFPruby::KanMX6_pTL58-pPma1-Ynl194c-5xGA-GFP::Leu2

Pil1-mRFPruby::KanMX6_pTL58-pPma1-Fmp45-5xGA-GFP::Leu2

Pil1-mRFPruby::KanMX6_pTL58-pPma1-Pun1-5xGA-GFP::Leu2

Pil1-mRFPruby::KanMX6_pTL58-pPma1-Sur7-5xGA-GFP::Leu2

Sur7-5xGA-mNeonGreen::hphNT1

Nce102-5xGA-mNeonGreen::hphNT1

Pil1-mRFPruby::KanMX6_pTL58-pPma1-Nce102-GAGA-GFP::Leu2

Sur7-HaloTag7::hphNT1

Nce102-HaloTag7::hphNT1

Pil1-mRFPruby::KanMX6_Tos7-5xGA-mNeonGreen::hphNT1

Mup1ΔC-HaloTag7::hphNT1

Pil1-mRFPruby::KanMX6_pTL58-pPma1-A08184g-5xGA-GFP::Leu2

pTL58-pPma1-Sur7-5xGA-HaloTag::Leu2

pTL58-pPma1-Ynl194c-5xGA-HaloTag::Leu2

pTL58-pPma1-Fmp45-5xGA-HaloTag::Leu2

pTL58-pPma1-Pun1-5xGA-HaloTag::Leu2

Δfmp45::_Δpun1::_Δsur7::_Δynl194::_Δtos7::

Δfmp45::_Δpun1::_Δsur7::_Δynl194::_Δtos7::_Nce102-5xGA-mNeonGreen::hphNT1

Δfmp45::_Δpun1::_Δsur7::_Δynl194::_Δtos7::_Nce102-HaloTag::hphNT1

Δfmp45::_Δpun1::_Δsur7::_Δynl194::_Δtos7::_Lsp1-HaloTag::hphNT1

Δfmp45::_Δpun1::_Δsur7::_Δynl194::_Δtos7::_Pil1-HaloTag::hphNT1

Lsp1-HaloTag::hphNT1

Pil1-HaloTag::hphNT1

Δfmp45::_Δpun1::_Δsur7::_Δynl194::_Δtos7:_Nce102-5xGA-mRFPRuby::natNT2_pTL58-pPma1-Sur7-5xGA-GFP::Leu2

Δfmp45::_Δpun1::_Δsur7::_Δynl194::_Δtos7:_Nce102-5xGA-mRFPRuby::natNT2_pTL58-pPma1-Ynl194c-5xGA-GFP::Leu2

Δfmp45::_Δpun1::_Δsur7::_Δynl194::_Δtos7:_Nce102-5xGA-mRFPRuby::natNT2_pTL58-pPma1-Fmp45-5xGA-GFP::Leu2

Δfmp45::_Δpun1::_Δsur7::_Δynl194::_Δtos7:_Nce102-5xGA-mRFPRuby::natNT2_pTL58-pPma1-Pun1-5xGA-GFP::Leu2

Δfmp45::_Δpun1::_Δsur7::_Δynl194::_Δtos7:_Nce102-5xGA-mRFPRuby::natNT2_pTL58-empty::Leu2

Δfmp45::_Δpun1::_Δsur7::_Δynl194::_Δtos7:_Nce102-5xGA-mRFPRuby::natNT2_pTL58-pPma1-A08184g-5xGA-GFP::Leu2

Lsp1-5xGA-mNeonGreen::hphNT1

Δfmp45::_Δpun1::_Δsur7::_Δynl194::_Δtos7:_Nce102-5xGA-mRFPRuby::natNT2_Inp51-5xGA-mNeonGreen::hphNT1

Nce102-5xGA-mRFPRuby::natNT2_Inp51-5xGA-mNeonGreen::hphNT1

BY4741:Sur7-5xGA-yeGFP_BY4742:Sur7-5xGA-mRFPRuby::Nat

Nce102-5xGA-mRFPruby::natNT2_pGPD-yeGFP-6xGA-Mss4::KanMX6

Pil1-5xGA-mRFPruby::KanMX6_Sur7-5xGA-mNeonGreen::hphNT1

Pil1-5xGA-mRFPruby::KanMX6_Pun1-5xGA-mNeonGreen::hphNT1

Pil1-5xGA-mRFPruby::KanMX6_Ynl194c-5xGA-mNeonGreen::hphNT1

Pil1-5xGA-mRFPruby::KanMX6_Nce102-5xGA-mNeonGreen::hphNT1

Pil1-5xGA-mRFPruby::KanMX6_Lsp1-5xGA-mNeonGreen::hphNT1

Δfmp45::_Δpun1::_Δsur7::_Δynl194::_Δtos7:_Nce102-5xGA-mRFPRuby::Nat_pGPD-yeGFP-6xGA-Mss4::KanMX6

Δfmp45::_Δpun1::_Δsur7::_Δynl194::_Δtos7:_Nce102-5xGA-mRFPRuby::natNT2_pTL58-pPma1-Tos7-5xGA-GFP::Leu2

Δfmp45::_Δpun1::_Δsur7::_Δynl194::_Δtos7:_Pil1-5xGA-mNeonGreen::natNT2

Pil1-5xGA-mNeonGreen::natNT2

Pil1-5xGA-mNeonGreen::natNT2_ Seg1-HaloTag::hphNT1

Δfmp45::_Δpun1::_Δsur7::_Δynl194::_Δtos7:_Pil1-5xGA-mNeonGreen::natNT2_Seg1-HaloTag::hphNT1

Pil1-5xGA-mNeonGreen::natNT2_ Sur7-HaloTag::hphNT1

Δinp51::natNT2_Δinp52::Ura3_Nce102-5xGA-mNeonGreen::hphNT1

Δinp51::natNT2_Δinp52::Ura3_Sur7-5xGA-mNeonGreen::hphNT1

Δfmp45::_Δpun1::_Δsur7::_Δynl194::_Δtos7:_Lsp1-5xGA-mNeonGreen::hphNT1

Δpsd1::natNT2_Δpsd2::Ura3_Sur7-5xGA-mNeonGreen::hphNT1

Δfmp45::_Δpun1::_Δsur7::_Δynl194::_Δtos7:_Nce102-HaloTag::hphNT1_pTL58-empty::Leu2

Δfmp45::_Δpun1::_Δsur7::_Δynl194::_Δtos7:_Δpil1::natNT2_Nce102-HaloTag::hphNT1

Pil1-GFP::HIS3MX6_pTL58-pPma1-Nce102-3xHA::Leu2

Sur7-GFP::HIS3MX6_pTL58-pPma1-Nce102-3xHA::Leu2

Δnce102::natNT2_Pil1-HaloTag::hphNT1

Δfmp45::_Δpun1::_Δsur7::_Δynl194::_Δtos7::_Δnce102::natNT2_Pil1-HaloTag::hphNT1

Nce102-5xGA-mRFPRuby::natNT2_pTL58-2xPH(PLCδ)-5xGA-GFP::Leu2

Δfmp45::_Δpun1::_Δsur7::_Δynl194::_Δtos7:_Nce102-5xGA-mRFPRuby::natNT2_pTL58-2xPH(PLCδ)-5xGA-GFP::Leu2

Δinp51::natNT2_Δinp52::Ura3_pTL58-Sur7-5xGA-HaloTag::Leu2

Δnce102::natNT2_pTL58-pPma1-Sur7-5xGA-HaloTag::Leu2

Δcho2::Ura3_Δopi3::natNT2_pTL58-pPma1-Sur7-5xGA-HaloTag::Leu2

Δcho1::natNT2_pTL58-pPma1-Sur7-5xGA-HaloTag::Leu2

Δpsd2::Ura3_Δpsd1::natNT2_pTL58-pPma1-Sur7-5xGA-HaloTag::Leu2

Δpil1::natNT2_Sur7-5xGA-HaloTag::hphNT1

Pil1-5xGA-HaloTag::hphNT1_pGPD-yeGFP-Seg1::KanMX6

Δfmp45::_Δpun1::_Δsur7::_Δynl194::_Δtos7::_Pil1-5xGA-HaloTag::hphNT1_pGPD-yeGFP-Seg1::KanMX6

Sur7-5xGA-HaloTag::hphNT1_pGPD-yeGFP-Seg1::KanMX6

Pil1-mNeonGreen::hphNT1_pTL58-pPma1-A08184g-5xGA-HaloTag::Leu2

pGPD-yeGFP-6xGA-Sur7::natNT2_pTL58-pPma1-Sur7-3xHA::Leu2

pGPD-yeGFP-6xGA-Fmp45::natNT2_pTL58-pPma1-Sur7-3xHA::Leu2

pGPD-yeGFP-6xGA-Ynl194c::natNT2_pTL58-pPma1-Sur7-3xHA::Leu2

pGPD-yeGFP-6xGA-Pun1::natNT2_pTL58-pPma1-Sur7-3xHA::Leu2

pGPD-yeGFP-6xGA-Seg1::KanMX6_pTL58-pPma1-Sur7-5xGA-HaloTag::Leu2

pTL58-pPma1-Tos7-HaloTag::Leu2

Δfmp45::_Δpun1::_Δsur7::_Δynl194::_Δtos7:_Nce102-5xGA-mRFPRuby::natNT2_pTL58-pPma1-Sur7(7-210)-5xGA-GFP::Leu2

Δfmp45::_Δpun1::_Δsur7::_Δynl194::_Δtos7:_Nce102-5xGA-mRFPRuby::natNT2_pTL58-pPma1-Sur7(7-302)-5xGA-GFP::Leu2

Δfmp45::_Δpun1::_Δsur7::_Δynl194::_Δtos7:_Nce102-5xGA-mRFPRuby::natNT2_pTL58-pPma1-Sur7(1-210)-5xGA-GFP::Leu2

pTL58-pPma1-Sur7(1-210)-5xGA-HaloTag::Leu2

Δfmp45::_Δpun1::_Δsur7::_Δynl194::_Δtos7_pTL58-pPma1-Sur7(1-210)-5xGA-HaloTag::Leu2

Nce102-5xGA-mRFPruby::natNT2_pTL58-pPma1-Sur7-5xGA-GFP::Leu2

Δfmp45::_Δpun1::_Δsur7::_Δynl194::_Δtos7_Δnce102::natNT2_Pil1-4A(S45A, S59A, S230A, T233A)-HaloTag::hphNT1

Pil1-HaloTag::hphNT1_pGPD-yeGFP-6xGA-Mss4::natNT2

Δfmp45::_Δpun1::_Δsur7::_Δynl194::_Δtos7_Pil1-HaloTag::hphNT1_pGPD-yeGFP-6xGA-Mss4::natNT2

Δfmp45::_Δpun1::_Δsur7::_Δynl194::_Δtos7::_Nce102-HaloTag::hphNT1_pTL58-pPma1-Sur7(7-210)-5xGA-GFP::Leu2

Δfmp45::_Δpun1::_Δsur7::_Δynl194::_Δtos7::_Nce102-HaloTag::hphNT1_pTL58-pPma1-Sur7(7-302)-5xGA-GFP::Leu2

Δfmp45::_Δpun1::_Δsur7::_Δynl194::_Δtos7::_Nce102-HaloTag::hphNT1_pTL58-pPma1-Sur7(1-210)-5xGA-GFP::Leu2

Δfmp45::_Δpun1::_Δsur7::_Δynl194::_Δtos7::_Nce102-HaloTag::hphNT1_pTL58-pPma1-Sur7-5xGA-GFP::Leu2

Pil1-5xGA-mRFPruby::KanMX6_pTL58-pPma1-Tos7-5xGA-GFP::Leu2

Pil1-5xGA-mRFPruby::KanMX6_Fmp45-5xGA-mNeonGreen::hphNT1

Δnce102::natNT2_Pil1-4A(S45A, S59A, S230A, T233A)-HaloTag::hphNT1

Pil1-5xGA-mNeonGreen::natNT2_Sur7-5xGA-mRFPruby::KanMX6

Δopi3::natNT2_Δcho2::Ura3_Sur7-5xGA-mNeonGreen::hphNT1

Δpil1::natNT2_Sur7-5xGA-mNeonGreen::hphNT1

Δcho1::natNT2_Sur7-5xGA-mNeonGreen::hphNT1

Nce102-3xHA::KanMX6_pTL58-pPma1-Nce102-5xGA-GFP::Leu2

Δfmp45::_Δpun1::_Δsur7::_Δynl194::_Δtos7::_Nce102-5xGA-mNeonGreen::hphNT1_Pil1-5xGA-mRFPruby::natNT2

Δfmp45::_Δpun1::_Δsur7::_Δynl194::_Δtos7::_Nce102-Sur7-chimera-5xGA-mNeonGreen::hphNT1

Δinp51::natNT2_Δinp52::Ura3_Pil1-HaloTag::hphNT1

Sur7-3xHA::KanMX6_pTL58-pPma1-Sur7(1-210)-5xGA-GFP::Leu2

Pil1-5xGA-mNeonGreen::hphNT1_pTL58-pPma1-Sur7-5xGA-HaloTag::Leu2

Sur7(1-210)-3xHA::KanMX6_pTL58-pPma1-Sur7-5xGA-GFP::Leu2

Sur7(1-210)-3xHA::KanMX6_pTL58-pPma1-Sur7(1-210)-5xGA-GFP::Leu2

Sur7-5xGA-mNeonGreen::hphNT1_Nce102-5xGA-mRFPruby::natNT2

Nce102-5xGA-mNeonGreen::hphNT1_Sur7-5xGA-mRFPruby::natNT2

Pil1-5xGA-mNeonGreen::hphNT1_Nce102-5xGA-mRFPruby::natNT2

Mup1ΔC-5xGA-mNeonGreen::hphNT1_Nce102-5xGA-mRFPruby::natNT2

Δinp51::natNT2_Δinp52::Ura3_Sur7-5xGA-mNeonGreen::hphNT1_Nce102-5xGA-mRFPruby::KanMX6

Δinp51::natNT2_Δinp52::Ura3_Nce102-Sur7-chimera-5xGA-mNeonGreen::hphNT1

Pil1-4A(S45A, S59A, S230A, T233A)-5xGA-mNeonGreen::hphNT1_Sur7-5xGA-mRFPruby::KanMX6

Δnce102::natNT2_Pil1-5xGA-mNeonGreen::hphNT1_Sur7-5xGA-mRFPruby::KanMX6

Δnce102::natNT2_Pil1-4A(S45A, S59A, S230A, T233A)-5xGA-mNeonGreen::hphNT1_Sur7-5xGA-mRFPruby::KanMX6

Δinp51::natNT2_Δinp52::Ura3_Nce102-5xGA-mNeonGreen::hphNT1_Sur7-5xGA-mRFPruby::KanMX6

Δinp51::natNT2_Δinp52::Ura3_Nce102-Sur7-chimera-5xGA-mNeonGreen::hphNT1_Sur7-5xGA-mRFPruby::KanMX6

| Source | Figure in Paper |
| --- | --- |
| Huh et al. 2003 |  |
| Euroscarf | Fig 9E |
| Euroscarf |  |
| this study | Fig 1F |
| this study | Fig 1F |
| this study | Fig 1F |
| this study | Fig 1F |
| this study | Fig 1A |
| this study | Fig 1A |
| this study | Fig 1A |
| this study | Fig 1A, Fig 6D |
| this study | Fig 2C; Fig 6A; Fig 9A; Fig 9D |
| this study | Fig 3G; Fig 3H; Fig EV3A; Fig 6E; Fig EV6A; Fig EV6B; Fig 9A; Fig 9D |
| this study | Fig 1A |
| this study | Fig 2A; Fig 2C; Fig 2D; Fig 7D |
| this study | Fig 2A; Fig 3D; Fig 4D |
| this study | Fig EV1A; Fig EV1E |
| this study | Fig 2A |
| this study | Fig 1A |
| this study | Fig 2B; Fig 2D; Fig EV2A; Fig 5C; Fig 6C; Fig 7D; Fig 8E; Fig 8G |
| this study | Fig EV2A |
| this study | Fig EV2A |
| this study | Fig EV2A |
| this study | Fig 9E |
| this study | Fig 3G; Fig 3H; Fig EV3A; Fig 6E; Fig EV6B; Fig 9A; Fig 9D |
| this study | Fig 3D; Fig 3F; Fig 4D |
| this study | Fig 3D |
| this study | Fig 3A; Fig 3F; Fig 5E; Fig 7A |
| this study | Fig 3D |
| this study | Fig 2A; Fig 2D; Fig 3A;F; Fig 5E; Fig 7A; Fig 8E |
| this study | Fig 4A; Fig 4C |
| this study | Fig 4A |
| this study | Fig 4A |
| this study | Fig 4A |
| this study | Fig 4A; Fig 4C |
| this study | Fig 4A |
| this study | Fig EV3A |
| this study | Fig 7E |
| this study | Fig 7E |
| this study | Fig 1D |
| this study | Fig 7F |
| this study | Fig EV1A; Fig EV1E; Fig 1D; Fig 8C; Fig 8D |
| this study | Fig EV1A; Fig EV1E |
| this study | Fig EV1A; Fig EV1E |
| this study | Fig 3E; Fig EV1A |
| this study | Fig 1D; Fig EV1E |
| this study | Fig 7F |
| this study | Fig 4A |
| this study | Fig EV3A |
| this study | Fig EV3A |
| this study | Fig 3D |
| this study | Fig 3D |
| this study | Fig EV2B |
| this study | Fig 9A; Fig 9D |
| this study | Fig 9D |
| this study | Fig EV3A |
| this study | Fig 6A |
| this study | Fig 4D |
| this study | Fig 3F; Fig 5E |
| this study | Fig 1F |
| this study | Fig 1F |
| this study | Fig 5E |
| this study | Fig 3F; Fig 5E |
| this study | Fig 7G; Fig 8A; Fig 8B |
| this study | Fig 7G; Fig 8B |
| this study | Fig 7D |
| this study | Fig 5C |
| this study | Fig 6C |
| this study | Fig 6C |
| this study | Fig 6C |
| this study | Fig 2C |
| this study | Fig 2D; Fig EV3B; Fig EV3D |
| this study | Fig EV3B; Fig EV3D |
| this study | Fig 2D |
| this study | Fig EV2A |
| this study | Fig EV1G |
| this study | Fig EV1G |
| this study | Fig EV1G |
| this study | Fig EV1G |
| this study | Fig 2D |
| this study | Fig EV2A |
| this study | Fig 4C |
| this study | Fig 4C |
| this study | Fig 4C |
| this study | Fig 4E |
| this study | Fig 4E |
| this study | Fig 4A; Fig 4C |
| this study | Fig 5E |
| this study | Fig 2D; Fig EV3B; Fig 7A |
| this study | Fig EV3B; Fig 7A |
| this study | Fig 4D |
| this study | Fig 4D |
| this study | Fig 4D |
| this study | Fig 4D |
| this study | Fig 1A |
| this study | Fig EV1A; Fig EV1E |
| this study | Fig 5E |
| this study | Fig 5A; Fig 5B |
| this study | Fig 6A |
| this study | Fig 2C |
| this study | Fig 6A |
| this study | Fig 1F |
| this study | Fig 3E |
| this study | Fig 9A; Fig 9D |
| this study | Fig 7A |
| this study | Fig 4F |
| this study | Fig 8F |
| this study | Fig 4F |
| this study | Fig 4F |
| this study | Fig 1D; Fig 7C; Fig EV1D |
| this study | Fig 9C |
| this study | Fig 1D |
| this study | Fig 1D |
| this study | Fig 7C |
| this study | Fig 9A |
| this study | Fig 5A; Fig 5B |
| this study | Fig 5A; Fig 5B |
| this study | Fig 5A; Fig 5B |
| this study | Fig 9C |
| this study | Fig 9C |

## comment

Background strain

Background strain

from *kluyveromyces lactis*

background strain for all 5xΔ cells

from *kluyveromyces lactis*

diploid yeast strain

background strains from Huh et al. 2003

background strains from Huh et al. 2003

